# Sustaining wakefulness: Brainstem connectivity in human consciousness

**DOI:** 10.1101/2023.07.13.548265

**Authors:** Brian L. Edlow, Mark Olchanyi, Holly J. Freeman, Jian Li, Chiara Maffei, Samuel B. Snider, Lilla Zöllei, J. Eugenio Iglesias, Jean Augustinack, Yelena G. Bodien, Robin L. Haynes, Douglas N. Greve, Bram R. Diamond, Allison Stevens, Joseph T. Giacino, Christophe Destrieux, Andre van der Kouwe, Emery N. Brown, Rebecca D. Folkerth, Bruce Fischl, Hannah C. Kinney

## Abstract

Consciousness is comprised of arousal (i.e., wakefulness) and awareness. Substantial progress has been made in mapping the cortical networks that modulate awareness in the human brain, but knowledge about the subcortical networks that sustain arousal is lacking. We integrated data from *ex vivo* diffusion MRI, immunohistochemistry, and *in vivo* 7 Tesla functional MRI to map the connectivity of a subcortical arousal network that we postulate sustains wakefulness in the resting, conscious human brain, analogous to the cortical default mode network (DMN) that is believed to sustain self-awareness. We identified nodes of the proposed default ascending arousal network (dAAN) in the brainstem, hypothalamus, thalamus, and basal forebrain by correlating *ex vivo* diffusion MRI with immunohistochemistry in three human brain specimens from neurologically normal individuals scanned at 600-750 µm resolution. We performed deterministic and probabilistic tractography analyses of the diffusion MRI data to map dAAN intra-network connections and dAAN-DMN internetwork connections. Using a newly developed network-based autopsy of the human brain that integrates *ex vivo* MRI and histopathology, we identified projection, association, and commissural pathways linking dAAN nodes with one another and with cortical DMN nodes, providing a structural architecture for the integration of arousal and awareness in human consciousness. We release the *ex vivo* diffusion MRI data, corresponding immunohistochemistry data, network-based autopsy methods, and a new brainstem dAAN atlas to support efforts to map the connectivity of human consciousness.

**One sentence summary:** We performed *ex vivo* diffusion MRI, immunohistochemistry, and *in vivo* 7 Tesla functional MRI to map brainstem connections that sustain wakefulness in human consciousness.

## INTRODUCTION

To treat disorders of human consciousness, such as coma, vegetative state, and minimally conscious state, it is essential to understand the neuroanatomic determinants and interconnectivity of arousal and awareness, two foundational components of consciousness (*1*). The goal of this multimodal brain mapping study is to advance knowledge of the connectivity of human consciousness by crystalizing the concept of a subcortical default arousal network that sustains wakefulness in the resting, conscious human brain, analogous to the cortical default mode network (DMN) that is believed to sustain self-awareness (*2*). We additionally provide a conceptual framework, imaging methods, and atlas-based reference tools that can be used to elucidate subcortical contributions to human consciousness.

Arousal refers to physiologic activation of the brain to a state of wakefulness (*3*), whereas awareness refers to the content of consciousness (*4*). These two components of consciousness – arousal and awareness – can be dissociated, as observed in patients in the vegetative state (*5*), who demonstrate alternating cycles of wakefulness but absent awareness (*5*). Such clinical observations, bolstered by experimental data in animals (*6-8*), support the concept that the anatomic pathways of awareness and arousal are anatomically dissociated from one another in the brain.

Current concepts of consciousness propose that the cerebral cortex is the primary anatomic site of the neural correlates of awareness (*4*), whereas ascending subcortical pathways from the brainstem, hypothalamus, thalamus, and basal forebrain are the neural correlates of arousal (*9, 10*). Over the past two decades, there has been substantial progress in mapping the cortical brain networks that mediate human awareness (*4, 11, 12*). In contrast, connectivity maps of subcortical networks that mediate human arousal are far less complete. This gap in knowledge is partly attributable to inadequate spatial resolution of standard neuroimaging techniques, which cannot discriminate individual arousal nuclei within the brainstem or delineate complex trajectories of axonal connections that link the brainstem to the diencephalon, basal forebrain, and cerebral cortex. In the absence of adequate imaging techniques for detecting brainstem arousal nuclei and their axonal connections, subcortical arousal network mapping has stalled while cortical awareness network mapping has accelerated.

The few human studies of brainstem arousal nuclei generally support Moruzzi and Magoun’s seminal observations in cats regarding the key role of the brainstem reticular formation in activating the cerebral cortex (*13*). Positron emission tomography (PET) studies have identified an increase in metabolic activity within the brainstem reticular formation when humans wake from sleep (*14*). Similarly, PET studies have revealed altered metabolism within the reticular formation in patients with disorders of consciousness caused by severe brain injuries (*15*).

Beyond the reticular formation, animal and human studies have revealed that extrareticular nuclei in the brainstem (*6, 16-19*), as well as nuclei in the hypothalamus (*7, 20, 21*), thalamus (*22, 23*), and basal forebrain (*6, 24, 25*), also contribute to arousal (*6, 26, 27*). Collectively, these observations suggest that the human ascending arousal network (AAN) is a synchronized, multi-transmitter brain network comprised of reticular and extrareticular nuclei that generate activated states of alertness in response to multiple external stimuli, including noxious, tactile, vestibular, olfactory, auditory, thermal, and homeostatic stimuli (e.g., hypoxia and hypercarbia). Our goal here is to identify the anatomic substrate of the AAN that modulates the *default state* of resting wakefulness, which we refer to as the default ascending arousal network (dAAN).

Default modes of brain activity involve neuronal ensembles that create distinct oscillatory modes of firing patterns during different brain states (*28*). In the awake state, the brain toggles between a default mode of wakefulness (i.e., resting wakefulness) and a non-default mode of wakefulness (i.e., active attention or task performance) (*2, 29*). In this conceptual framework, a default network is one whose nodes demonstrate temporally correlated activity during a default mode of brain activity (*30*). Maintenance of the resting wakeful state requires input from the dAAN. This view of a default state of arousal is consistent with that expressed by Steriade and Buzsaki, who wrote that “operationally, arousal may be defined as a stand-by mode of the neurons for efficient processing and transformation of afferent information” (*31*). In the “stand-by mode” of resting wakefulness, the human brain is not performing an externally or internally directed (i.e., introspective) task. Instead, the mind wanders and stimulus-independent thoughts may emerge (*32*). Behaviorally, the eyes may be open or closed. Electrophysiologically, subcortical arousal neurons fire tonically (*33*) and the cerebral cortex generates desynchronized, high frequency, low voltage waves on EEG (*3*). The dAAN thus sustains cortical activation to enable awareness, without requiring sensory input or externally directed cognition.

To advance knowledge about the connectivity of this subcortical default brain network critical to human consciousness, we performed a multimodal brain imaging study, integrating data from *ex vivo* diffusion MRI tractography and *in vivo* 7 Tesla functional MRI. We used histology and immunohistochemistry guidance to identify the anatomic location of dAAN nodes on *ex vivo* MRI scans of three human brain specimens, then mapped the structural connections of the dAAN using diffusion MRI tractography at submillimeter spatial resolution. We then performed resting-state functional MRI (rs-fMRI) analyses of 7 Tesla imaging data from the Human Connectome Project (*34*) to investigate the functional correlates of these structural connections. The complementary structural-functional connectivity analyses aimed to identify the neuroanatomic basis for the integration of arousal and awareness in the resting, conscious human brain. We release all network-based autopsy methods and data, and we propose a neuroanatomic taxonomy for projection, association, and commissural pathways of the human dAAN to guide future investigations into the connectivity of human consciousness and its disorders.

## RESULTS

### Candidate dAAN nodes

Based on evidence from previously published electrophysiological, gene expression, lesion, and stimulation studies, we identified 18 candidate dAAN nodes: 10 brainstem, 3 thalamus, 3 hypothalamus, and 2 basal forebrain (**Table 1**). All 18 candidate nodes were located at the level of, or rostral to, the mid-pons. For the 13 nodes with available electrophysiological data (all except periaqueductal grey [PAG], pontis oralis [PnO], parabrachial complex [PBC], paraventricular nucleus [PaV], and supramammillary nucleus [SuM]), the electrophysiological evidence indicated active tonic firing of neurons during wakefulness and state-dependent changes in neuronal firing during wakefulness as compared to slow-wave sleep (**Table 1**).

**Table 1.**
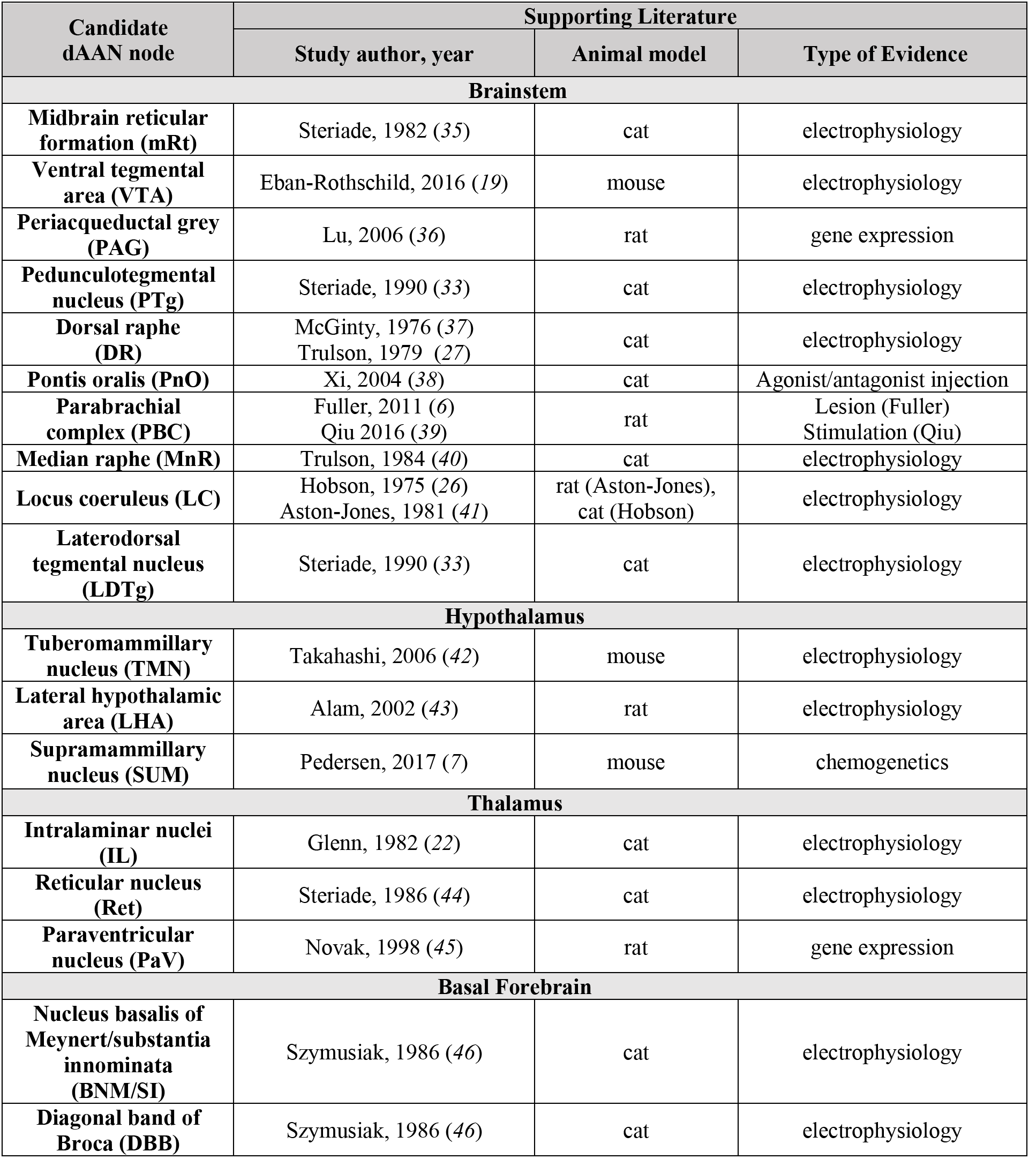
Candidate default ascending arousal network (dAAN) nodes. We identified 18 nodes that met the *a priori* criteria for inclusion in the human dAAN. The criteria were based on previous animal studies demonstrating nodal involvement in arousal from electrophysiological recording, Fos expression, lesioning, or stimulation. Given that decades of experiments in multiple animal species have tested the functional properties of subcortical arousal nuclei, we do not intend for the references in this table to be exhaustive. Rather, we list studies that are representative of the body of literature pertaining to each subcortical arousal nucleus.

| Candidate dAAN node | Supporting Literature |  |  |
| --- | --- | --- | --- |
|  | Study author, year | Animal model | Type of Evidence |
| **Brainstem** |  |  |  |
| **Midbrain reticular formation (mRt)** | Steriade, 1982 (35) | cat | electrophysiology |
| **Ventral tegmental area (VTA)** | Eban-Rothschild, 2016 (19) | mouse | electrophysiology |
| **Periaqueductal grey (PAG)** | Lu, 2006 (36) | rat | gene expression |
| **Pedunculotegmental nucleus (PTg)** | Steriade, 1990 (33) | cat | electrophysiology |
| **Dorsal raphe (DR)** | McGinty, 1976 (37)<br>Trulson, 1979 (27) | cat | electrophysiology |
| **Pontis oralis (PnO)** | Xi, 2004 (38) | cat | Agonist/antagonist injection |
| **Parabrachial complex (PBC)** | Fuller, 2011 (6)<br>Qiu 2016 (39) | rat | Lesion (Fuller)<br>Stimulation (Qiu) |
| **Median raphe (MnR)** | Trulson, 1984 (40) | cat | electrophysiology |
| **Locus coeruleus (LC)** | Hobson, 1975 (26)<br>Aston-Jones, 1981 (41) | rat (Aston-Jones),<br>cat (Hobson) | electrophysiology |
| **Laterodorsal tegmental nucleus (LDTg)** | Steriade, 1990 (33) | cat | electrophysiology |
| **Hypothalamus** |  |  |  |
| **Tuberomammillary nucleus (TMN)** | Takahashi, 2006 (42) | mouse | electrophysiology |
| **Lateral hypothalamic area (LHA)** | Alam, 2002 (43) | rat | electrophysiology |
| **Supramammillary nucleus (SUM)** | Pedersen, 2017 (7) | mouse | chemogenetics |
| **Thalamus** |  |  |  |
| **Intralaminar nuclei (IL)** | Glenn, 1982 (22) | cat | electrophysiology |
| **Reticular nucleus (Ret)** | Steriade, 1986 (44) | cat | electrophysiology |
| **Paraventricular nucleus (PaV)** | Novak, 1998 (45) | rat | gene expression |
| **Basal Forebrain** |  |  |  |
| **Nucleus basalis of Meynert/substantia innominata (BNM/SI)** | Szymusiak, 1986 (46) | cat | electrophysiology |
| **Diagonal band of Broca (DBB)** | Szymusiak, 1986 (46) | cat | electrophysiology |

### dAAN structural connectivity

#### Human brain specimens

Once the 18 candidate dAAN nodes were identified, we investigated their structural connectivity using *ex vivo* diffusion MRI tractography in three human brain specimens from neurologically normal individuals. The three brain specimens were from women who died at ages 53, 60, and 61 years (**Fig. S1-S3**). Demographic and clinical data, including cause of death, are provided in **Table S1**. At autopsy, all three brain specimens had a normal fresh brain weight. Time from death until fixation in 10% formaldehyde ranged from 24 hours to 72 hours, a time window that is associated with preservation of directional water diffusion within formaldehyde-fixed brain tissue (*47*).

For specimen 1, candidate dAAN nodes were traced on the diffusion-weighted images using guidance from serial histologic sections (hematoxylin and eosin counterstained with Luxol fast blue) and selected immunohistochemistry sections (tyrosine hydroxylase for dopaminergic neurons and tryptophan hydroxylase for serotonergic neurons). Each set of diffusion images (diffusion-weighted images [DWI], apparent diffusion coefficient [ADC] map, fractional anisotropy [FA] map, color FA map, and non-diffusion weighted images [*b=0*]) was compared to its corresponding stained sections to ensure that radiologic dAAN nodes shared the same location and morphology as the dAAN nuclei identified by histology and immunohistochemistry (**Figs. 1, S4, S5**). For specimens 2 and 3, the brainstem dAAN nuclei were traced manually on each diffusion MRI dataset with reference to selected histologic and immunohistochemistry sections and canonical atlases (*48, 49*). All histology and immunostaining data are available at https://histopath.nmr.mgh.harvard.edu.

**Fig. 1.**
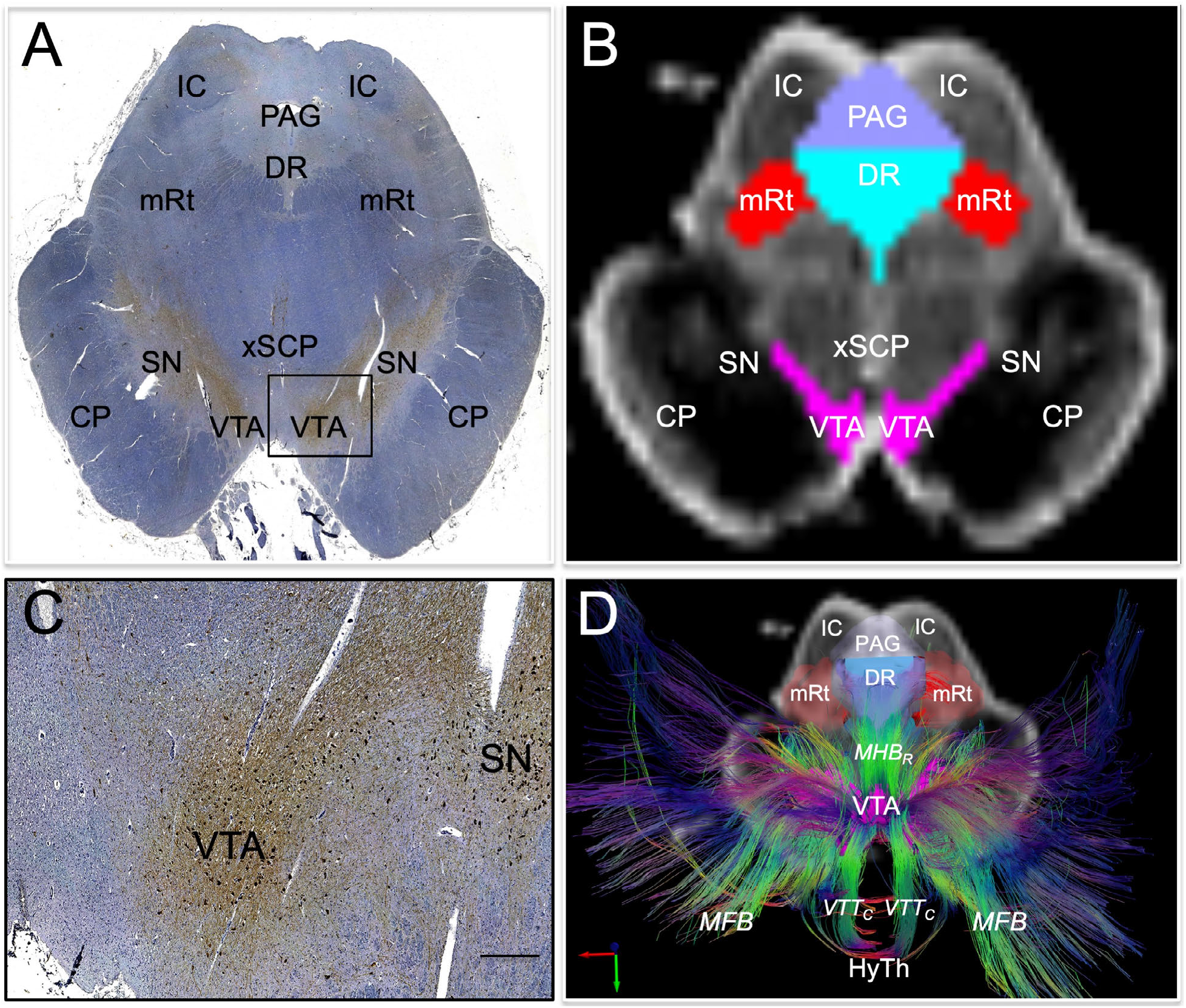
Histological guidance of node localization and tract construction. **(A)** A transverse section through the caudal midbrain of specimen 1 is shown, stained with tyrosine hydroxylase and counterstained with hematoxylin. The tyrosine hydroxylase stain identifies dopaminergic neurons of the ventral tegmental area (VTA). A zoomed view of VTA neurons located within the rectangle in **(A)** are shown in **(C;** scale bar = 100 µm). The corresponding non-diffusion-weighted (*b=0*) axial image from the same specimen is shown in **(B)**. VTA neurons are manually traced in pink, based upon the tyrosine hydroxylase staining results, and additional arousal nuclei are traced based on the hematoxylin and eosin/Luxol fast blue stain results: DR = dorsal raphe; mRt = mesencephalic reticular formation; PAG = periaqueductal grey. Deterministic fiber tracts emanating from the VTA node are shown in **(D)** from the same superior view as panel **(B)**. The tracts are color-coded by direction (inset, bottom left) and the DR and mRt nodes are semi-transparent so that VTA tracts can be seen passing through them via the mesencephalic homeostatic bundle, rostral division (MHBR). VTA tracts also connect with the hypothalamus (HyTh), thalamus, basal forebrain, and cerebral cortex via multiple bundles: MFB = medial forebrain bundle; VTT_C_ = ventral tegmental tract, caudal division. Additional abbreviations: CP = cerebral peduncle; IC = inferior colliculus; SN = substantia nigra; xSCP = decussation of the superior cerebellar peduncles.

To facilitate reproduction of our methods, we detail the neuroanatomic boundaries of all candidate dAAN nodes in **Tables S2-S6**. In addition, we provide a comprehensive summary of our anatomic approach in the **Supplementary Methods**, including a discussion of how the neuroanatomic borders of our candidate dAAN nodes relate to those defined in canonical atlases (*50-52*) and how they differ from those of the nodes distributed in version 1.0 of the Harvard Ascending Arousal Network Atlas (*9*). Based on the updated neuroanatomic nomenclature and nodal anatomic characteristics reported here, we release version 2.0 of the Harvard Ascending Arousal Network Atlas (**Video S1**) as a tool for the academic community (doi:10.5061/dryad.zw3r228d2; download link: https://datadryad.org/stash/share/aJ713eXY12ND56bzOBejVG2jmOFCD2CKxdSJsYWEHkw).

Specimen 1 was scanned at 600 µm resolution as a dissected specimen, consisting of the rostral pons, midbrain, hypothalamus, thalamus, basal forebrain, and part of the basal ganglia (*9*). Specimens 2 and 3 were scanned at 750 µm resolution as whole brains. Diffusion data from all three specimens were processed for deterministic fiber tracking for qualitative assessment of tract trajectories, and diffusion data from specimens 2 and 3 were processed for probabilistic fiber tracking for quantitative assessment of connectivity. Probabilistic data for Specimen 1 were not analyzed because differences in scanning parameters and specimen composition – (i.e., dissected versus whole brain) precluded quantitative comparison. We performed deterministic tractography to assess qualitative anatomic relationships of dAAN tracts, and probabilistic tractography to measure quantitative dAAN connectivity properties.

#### Criteria for structural connectivity and anatomic classification of dAAN pathways

Structural connectivity was defined using a two-step process that incorporated the quantitative probabilistic data and qualitative deterministic data. First, we performed a quantitative assessment for the presence of a connection linking each node-node pair using the probabilistic data for specimens 2 and 3 (see statistical analysis section of the Materials and Methods). A significant quantitative connection requires a connectivity probability measure that exceeds the 95% confidence interval of connectivity with two control regions that are unlikely to have connectivity with dAAN nodes based on prior animal labeling studies (e.g., (*53*)): basis pontis and red nucleus. If this *a priori* criterion was met in the probabilistic analysis, we then qualitatively assessed the deterministic tractography data for all three specimens for visual confirmation of tracts connecting each pair of candidate nodes. Node-node pairs that met both the probabilistic and deterministic criteria were considered to have inferential evidence for a structural connection, while node-node pairs that met one, but not both, criteria were considered to have uncertain connectivity.

We then classified the structural connectivity between all candidate node-node pairs based on the anatomic trajectories and nomenclature of subcortical bundles previously defined in humans and animals (*9, 54*): DTT_L_ = dorsal tegmental tract, lateral division; DTT_M_ = dorsal tegmental tract, medial division; LFB = lateral forebrain bundle; MFB = medial forebrain bundle; VTT_C_ = ventral tegmental tract, caudal division; VTT_R_ = ventral tegmental tract, rostral division (**Fig. 2**). Next, we classified all dAAN pathways as projection, association, or commissural, providing a neuroanatomic taxonomy for the subcortical dAAN, akin to that of cortical networks whose connections are classified in these terms (*55*). Using the brainstem as the frame of reference for classifying dAAN pathways, we defined projection fibers as those that project from brainstem nuclei to rostral diencephalic nuclei, basal forebrain nuclei, and/or cortical regions that mediate consciousness (i.e., DMN nodes). Association fibers connect brainstem nuclei to ipsilateral brainstem nuclei, and commissural fibers cross the midline to connect with contralateral brainstem nuclei.

**Fig. 2.**
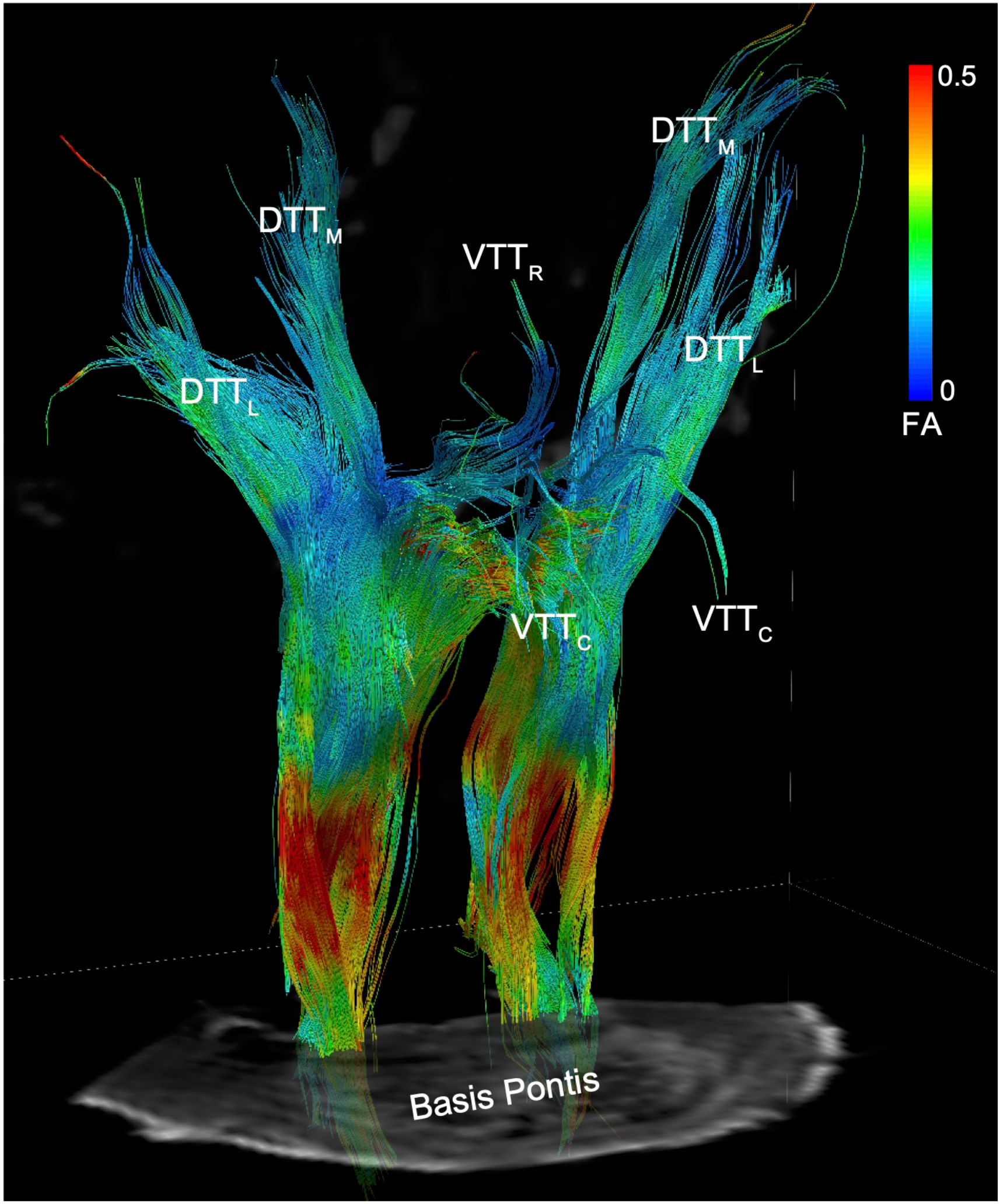
Reticular pathways of the human default ascending arousal network. Deterministic tractography results from a seed region in the mesencephalic reticular formation (mRt), the brainstem region stimulated by Moruzzi and Magoun in their seminal investigations of the reticular activating system (*13*), are shown from a right ventro-lateral perspective in Specimen 1. Tracts are color-coded by the fractional anisotropy (FA) along each segment (inset color bar). Tracts are superimposed upon a non-diffusion-weighted (*b=0*) axial image at the level of the mid-pons. The fiber tracts emanating from mRt travel in the ponto-mesencephalic tegmentum and connect with the thalamus, hypothalamus, and basal forebrain via the following bundles: DTT_L_ = dorsal tegmental tract, lateral division; DTT_M_ = dorsal tegmental tract, medial division; VTT_C_ = ventral tegmental tract, caudal division; VTT_R_ = ventral tegmental tract, rostral division.

#### Projection pathways of the dAAN

Deterministic tractography analysis of the three specimens revealed that tracts emanating from all dAAN candidate brainstem nuclei formed well-defined projection pathways that coursed through the rostral brainstem, diencephalon, and forebrain. Each pathway contained intermingled tracts from reticular nuclei (i.e., pontine and mesencephalic reticular formations) and extrareticular nuclei (e.g., monoaminergic, cholinergic, and glutamatergic nuclei), though the distributions of reticular and extrareticular tracts within each pathway differed (**Fig. 3**).

**Fig. 3.**
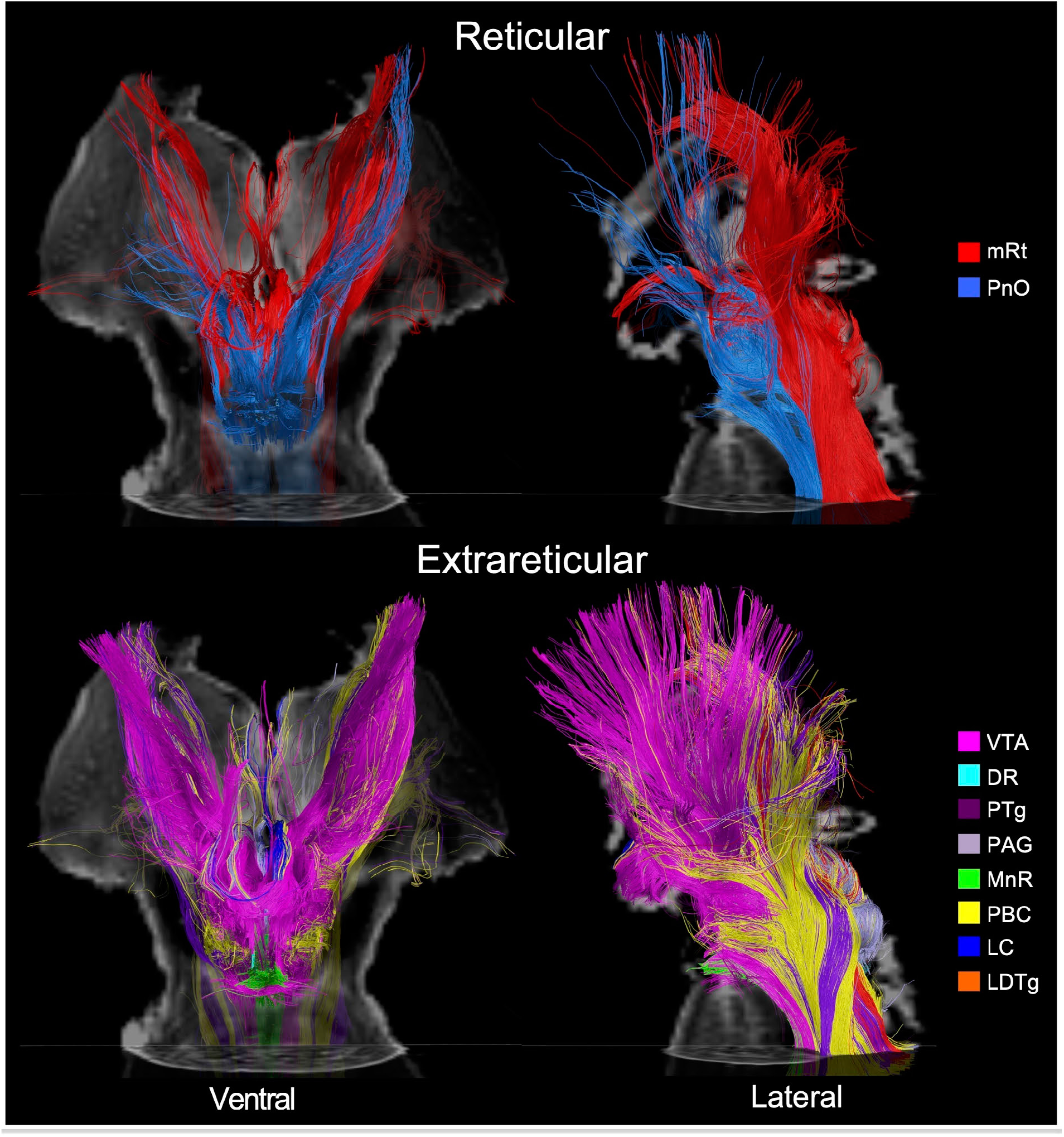
Reticular and extrareticular tracts of the default ascending arousal network. Deterministic fiber tracts emanating from the reticular arousal nuclei are shown in the top panels. Fiber tracts emanating from extrareticular arousal nuclei are shown in the bottom panels. All tracts are from specimen 1 and superimposed on an axial non-diffusion-weighted (*b=0*) image at the level of the mid-pons. Additionally, the tracts in the ventral perspective (left column) are superimposed on a coronal *b=0* image at the level of the mid-thalamus, and the tracts in the left lateral perspective (right column) are superimposed on a sagittal *b=0* image located at the midline. All tracts are color-coded according to their nucleus of origin (inset, right). Abbreviations are described in Table 1.

All candidate brainstem nodes connected with at least one candidate hypothalamic, thalamic, or basal forebrain node based on confidence interval testing (**Table 2**). Quantitative connectivity data are provided in **Tables S7-S10** for all projection pathways. The projection pathways that connected candidate brainstem dAAN nodes with thalamic, hypothalamic, and basal forebrain dAAN nodes are shown in **Fig. 3**. The projection pathways connecting candidate brainstem dAAN nodes with cortical DMN nodes are shown in **Fig. 4**. A projection fiber connectogram with averaged CP values for specimens 2 and 3 is shown in **Fig. 5**.

**Table 2.** Projection pathways connecting brainstem dAAN nodes with thalamic, hypothalamic, and basal forebrain dAAN nodes, as well as cortical DMN nodes. For connections that met both the probabilistic and deterministic criteria for a structural connection, we assessed the deterministic fiber tracts linking each pair of nodes for their anatomic trajectories and classified them using previously defined nomenclature (*9, 54*): DTT_L_ = dorsal tegmental tract, lateral division; DTT_M_ = dorsal tegmental tract, medial division; LFB = lateral forebrain bundle; MFB = medial forebrain bundle; VTT_C_ = ventral tegmental tract, caudal division; VTT_R_ = ventral tegmental tract, rostral division. If two nodes did not meet the criteria for a connection using the probabilistic or determinist criteria, the label “---” is given, to indicate the absence of a connection. If two nodes met criteria for a connection in the probabilistic or deterministic analyses, but not both analyses, then the label “UC” (uncertain) is given. * The DTT_L_ pathway connects directly with Ret, whereas the DTT_M_ pathway connects with Ret after passing through IL.

| Brainstem Node | Thalamus |  |  | Hypothalamus |  |  | Basal Forebrain |  | Default Mode Network |  |  |  |  |
| --- | --- | --- | --- | --- | --- | --- | --- | --- | --- | --- | --- | --- | --- |
|  | IL | Ret* | PaV | TMN | LHA | SUM | BNM/SI | DBB | PMC | MPFC | IPL | LT | MT |
| **mRt** | DTT<sub>M</sub> | DTT<sub>L</sub><br>DTT<sub>M</sub> | VTT<sub>R</sub> | VTT<sub>C</sub> | VTT<sub>C</sub> | VTT<sub>C</sub> | VTT<sub>C</sub> | VTT<sub>C</sub> | LFB | MFB | LFB | UC | LFB |
| **VTA** | DTT<sub>M</sub> | UC | VTT<sub>R</sub> | VTT<sub>C</sub> | VTT<sub>C</sub> | VTT<sub>C</sub> | VTT<sub>C</sub> | UC | LFB | MFB | LFB | LFB | LFB |
| **PAG** | DTT<sub>M</sub> | DTT<sub>L</sub><br>DTT<sub>M</sub> | VTT<sub>R</sub> | --- | VTT<sub>C</sub> | VTT<sub>C</sub> | VTT<sub>C</sub> | VTT<sub>C</sub> | LFB | UC | LFB | LFB | LFB |
| **PTg** | DTT<sub>M</sub> | DTT<sub>L</sub><br>DTT<sub>M</sub> | VTT<sub>R</sub> | VTT<sub>C</sub> | VTT<sub>C</sub> | VTT<sub>C</sub> | VTT<sub>C</sub> | VTT<sub>C</sub> | LFB | MFB | UC | LFB | LFB |
| **DR** | DTT<sub>M</sub> | DTT<sub>L</sub> | VTT<sub>R</sub> | VTT<sub>C</sub> | VTT<sub>C</sub> | VTT<sub>C</sub> | VTT<sub>C</sub> | VTT<sub>C</sub> | LFB | UC | UC | UC | LFB |
| **PnO** | DTT<sub>M</sub> | DTT<sub>L</sub><br>DTT<sub>M</sub> | --- | --- | VTT<sub>C</sub> | --- | VTT<sub>C</sub> | --- | LFB | MFB | UC | UC | --- |
| **PBC** | DTT<sub>M</sub> | DTT<sub>L</sub><br>DTT<sub>M</sub> | UC | VTT<sub>C</sub> | VTT<sub>C</sub> | --- | VTT<sub>C</sub> | UC | LFB | MFB | LFB | UC | UC |
| **MnR** | --- | --- | --- | --- | VTT<sub>C</sub> | --- | VTT<sub>C</sub> | --- | LFB | MFB | LFB | UC | --- |
| **LC** | UC | UC | UC | VTT<sub>C</sub> | VTT<sub>C</sub> | VTT<sub>C</sub> | VTT<sub>C</sub> | UC | LFB | UC | UC | UC | UC |
| **LDTg** | UC | UC | --- | VTT<sub>C</sub> | VTT<sub>C</sub> | VTT<sub>C</sub> | UC | VTT<sub>C</sub> | UC | UC | UC | UC | UC |

**Fig. 4.**
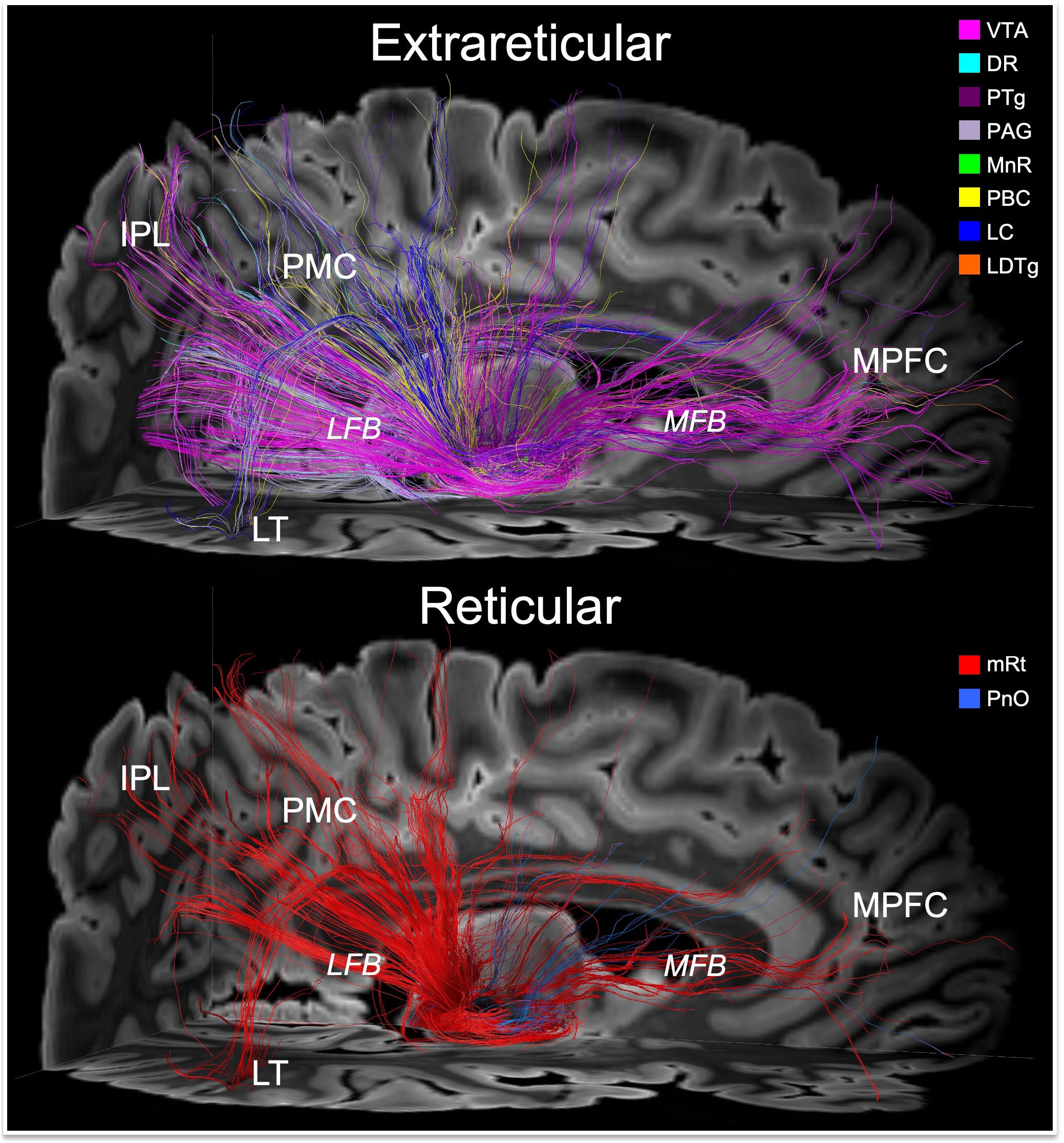
Connections linking brainstem nodes of the default ascending arousal network with cortical nodes of the default mode network. Deterministic fiber tracts emanating from the extrareticular and reticular arousal nuclei are shown in the top and bottom panels, respectively. All tracts are shown from a right lateral perspective and superimposed upon an axial non-diffusion-weighted (*b=0*) image at the level of the rostral midbrain and a sagittal *b=0* image to the right of the midline. Tract colors and abbreviations are the same as in Fig. 3. Both the extrareticular and reticular brainstem arousal nuclei connect extensively with cortical nodes of the default mode network, including the medial prefrontal cortex (MPFC), posteromedial complex (PMC; i.e., posterior cingulate and precuneus), inferior parietal lobule (IPL), and lateral temporal lobe (LT). The primary pathway that connects the brainstem arousal nuclei to MPFC is the medial forebrain bundle (MFB). The primary pathway that connects the brainstem arousal nuclei to the PMC, IPL and LT is the lateral forebrain bundle (LFB).

**Fig. 5.**
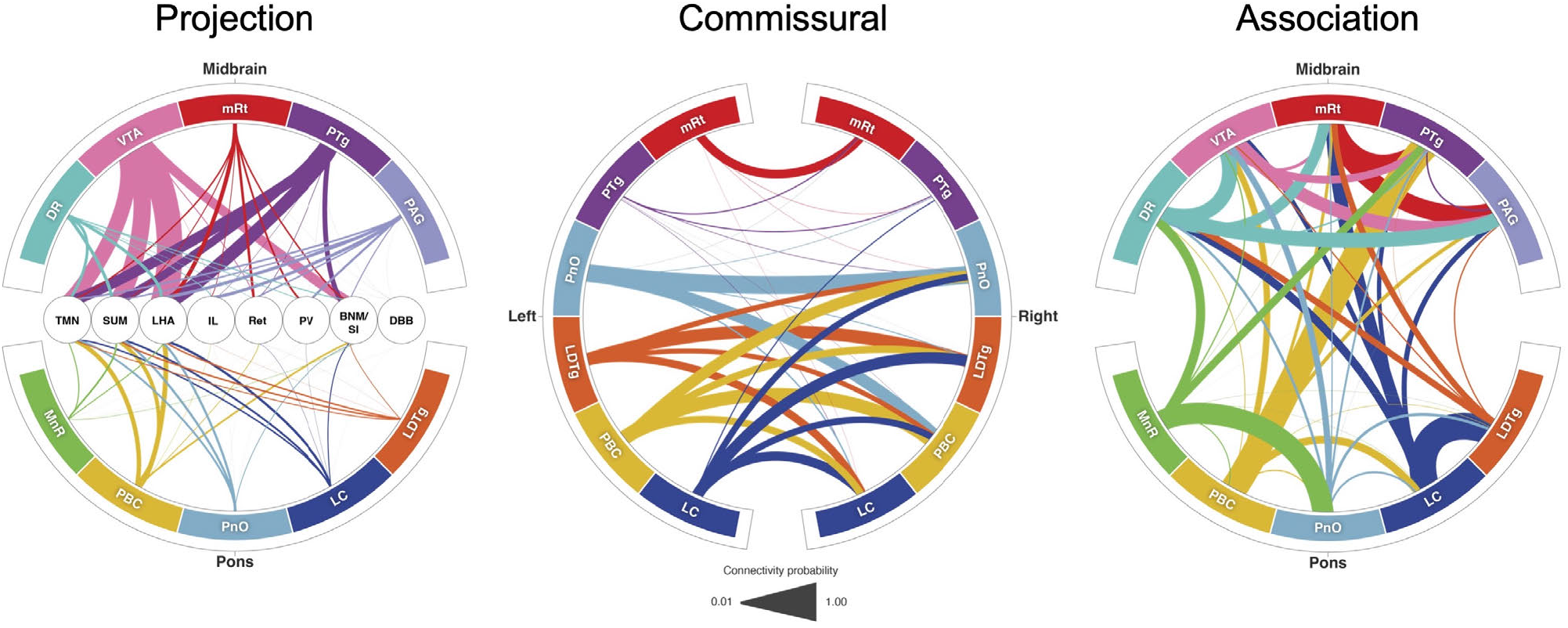
Default ascending arousal network connectograms. Projection pathways (left) connect brainstem nodes to hypothalamic, thalamic, and basal forebrain nodes. Commissural pathways (middle) connect contralateral brainstem nodes. Association pathways (right) connect ipsilateral and/or midline brainstem nodes. Brainstem dAAN nodes are shown in the outer ring of all connectograms. In the projection connectogram, the hypothalamic, thalamic and basal forebrain nodes are shown in the center: TMN = tuberomammillary nucleus; LHA = lateral hypothalamic area; IL = intralaminar nuclei; Ret = reticular nuclei; PaV = paraventricular nuclei; BNM/SI = nucleus basalis of Meynert/substantia innominata; DBB = diagonal band of Broca. Line thicknesses in each connectogram is proportional to the connectivity probability (CP) value derived from probabilistic tractography analysis. For clarity of visualization, interspecimen mean in CP measurements is shown in the connectograms. In the commissural connectogram, we show the connections from left-to-right and omit the right-to-left connections for ease of visualization. In the association connectogram, we take a mean of left- and right-sided connectivity for bilaterally represented nodes. We do not show a connectogram for projection pathways to the DMN nodes because the connectivity probability values are lower than for diencephalic and forebrain projection pathways (Table S10), such that connections would not be visualizable at this scale.

The qualitative trajectories of projection pathways connecting these nodes, as assessed by deterministic tractography, were consistent in all three specimens. All midbrain candidate nodes connected with IL via DTT_M_ and with PaV via VTT_R_. All midbrain nodes except VTA (which showed uncertain connectivity) connected with Ret via DTT_L_. Connectivity between pontine candidate nodes and the thalamic candidate nodes was more limited, with only PnO and PBC connecting with IL and Ret, and with MnR, LC, and LDTg showing either absent or uncertain connectivity with thalamic nodes (see Discussion for potential explanations of the latter finding).

Structural connectivity with all three candidate hypothalamic nodes (TMN, LHA, and SUM) via VTT_C_ was observed for mRt, VTA, PTg, DR, LC, and LDTg, whereas other brainstem nuclei connected with one (PnO, MnR) or two (PAG, PBC) candidate hypothalamic nodes. All midbrain and pontine candidate nodes connected with the LHA node of the hypothalamus via VTT_C_. Structural connectivity with the NBM/SI node of the basal forebrain was observed for all midbrain and pontine candidate nodes except LDTg via VTT_C_. All brainstem nodes except VTA, PnO, PBC, MnR, and LC connected with the DBB node of the basal forebrain via VTT_C_.

All cortical DMN nodes showed structural connectivity with at least 3 of the 10 candidate brainstem dAAN nodes. The PMC (i.e., posterior cingulate and precuneus) was the DMN node with the most extensive connections to brainstem dAAN nodes (all except LDTg), via the LFB. The MPFC node of the DMN was connected with all brainstem dAAN nodes except PAG, DR, LC and LDTg, via the MFB. The IPL node of the DMN connected with mRt, VTA, PAG, PBC, and MnR via LFB. The MT node of the DMN was connected with all midbrain nodes via a temporal branch of the LFB, and the LT node of the DMN was connected with all midbrain nodes except mRt. There were no connections between LT or MT and the pontine candidate dAAN nodes. We observed a large contribution of extrareticular connections, dominated by VTA tracts, to the MPFC via the MFB (**Fig. 4**). Brainstem connections with IPL and PMC traveled in a parietal branch of the LFB, with similar connectivity contributions from VTA, PAG and mRt (**Fig. 4**).

#### Association pathways of the dAAN

Deterministic and probabilistic tractography analyses revealed connections linking all candidate brainstem nodes with at least one ipsilateral or midline brainstem node (**Fig. 5**). As with the projection fiber pathways, the association fiber pathways were similar between specimens. Extensive connectivity between midline, left-sided, and right-sided brainstem candidate dAAN nodes (both reticular and extrareticular) was observed (**Fig. 6** and **Video S2**). Quantitative connectivity data from the probabilistic tractography analyses of association pathways are provided in **Tables S11, S12, and S13**.

**Fig. 6.**
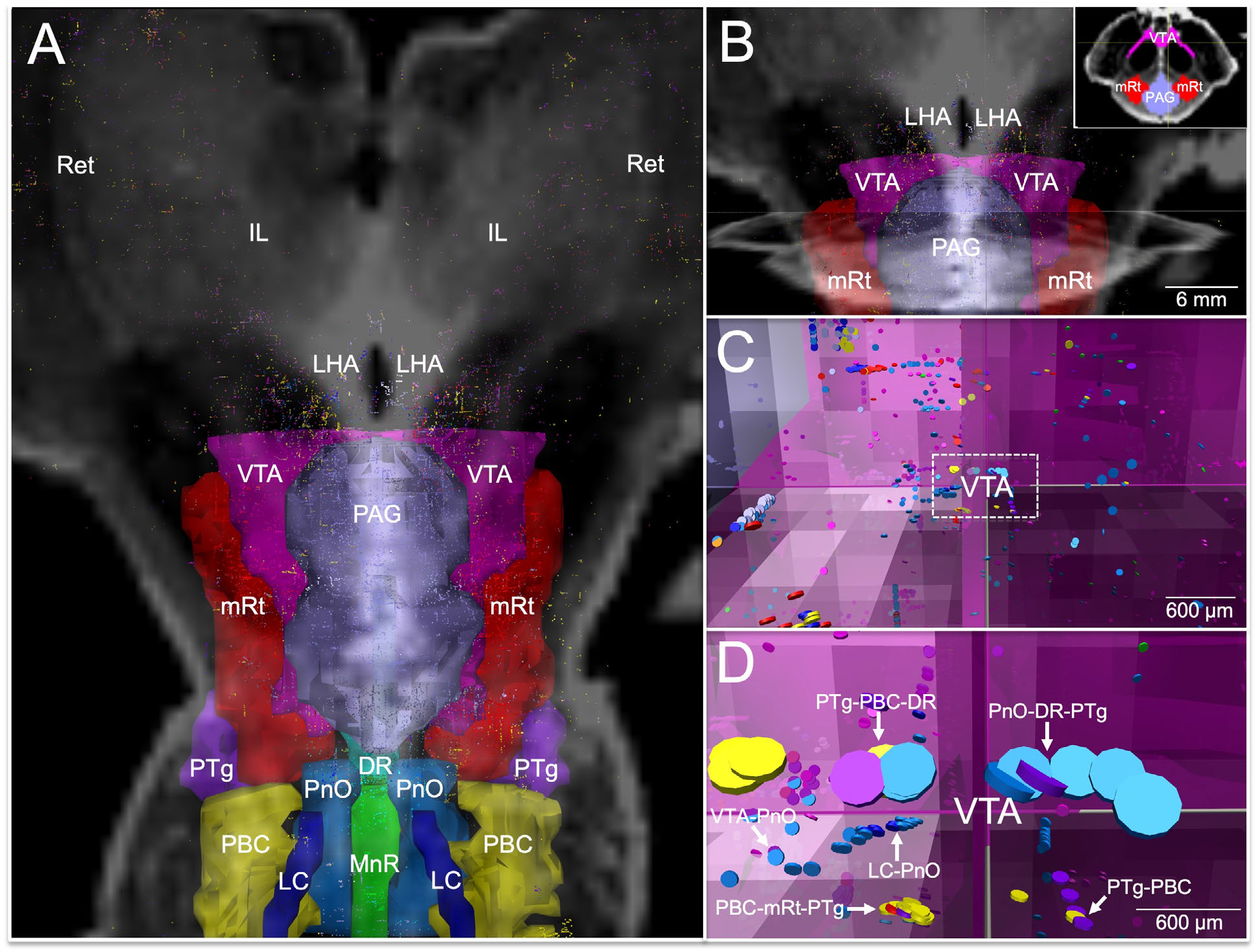
Connections linking association pathways of the default arousal network. All candidate brainstem dAAN nodes and their tract end-points are shown from a dorsal perspective in **(A)**, superimposed upon a coronal non-diffusion-weighted (*b=0*) image at the level of the mid-thalamus. Tract end-points represent the start-points and termination-points of each tract (i.e., there are two end-points per tract). All tract end-points are color-coded by the node from which they originate, and the nodes are rendered semi-transparent so that end-points can be seen within them. In **(A)**, tract end-points are seen within each brainstem candidate node, as well as in the lateral hypothalamic area (LHA), intralaminar nuclei of the thalamus (IL), and reticular nuclei of the thalamus (Ret). A zoomed-in view of the tract end-points within the ventral tegmental area (VTA), periaqueductal grey (PAG), and mesencephalic reticular formation (mRt) is shown in **(B**), superimposed upon the same coronal *b=0* image as in **(A)**, as well as an axial *b=0* image at the intercollicular level of the midbrain. Low-zoom and high-zoom views of tract end-points (appearing as discs) within VTA are shown in **(C)** and **(D)**, respectively; the axes shown in **(B, C,** and **D)** are identical and are located at the dorsomedial border of the VTA node, as shown in the **Panel B** inset. In **(C** and **D),** the perspective is now ventral to the PAG, just at the dorsomedial border of the VTA, as indicated by the axes. Tract end-points from multiple candidate brainstem dAAN nodes are seen overlapping within the VTA and along its dorsal border (arrows): pedunculotegmental, parabrachial complex, and dorsal raphe (PTg-PBC-DR); pontis oralis, DR, and PTg (PnO-DR-PTg); VTA-PnO; PBC-mRt-PTg; locus coeruleus and PnO (LC-PnO); and PTg-PBC. Tract end-point overlap implies that that the beginning and/or endpoints of two tracts are in very close proximity, suggesting extensive connectivity via association pathways between ipsilateral and midline dAAN nodes with the VTA.

#### Commissural pathways of the dAAN

We observed fewer connections in the commissural analyses as compared to the projection or association analyses. Whereas projection and association tracts were distributed in multiple discreet pathways (e.g., DTT_M_, DTT_L_, VTT_R_, VTT_C_), the commissural tracts were mostly seen within a single pathway traveling across the posterior commissure in the midbrain (**Fig. 7**). Additional commissural tracts outside the posterior commissure, particularly those in the pons, did not coalesce into discrete bundles of tracts. Quantitative connectivity data from the probabilistic tractography analyses of commissural pathways are provided in **Table S14**.

**Fig. 7.**
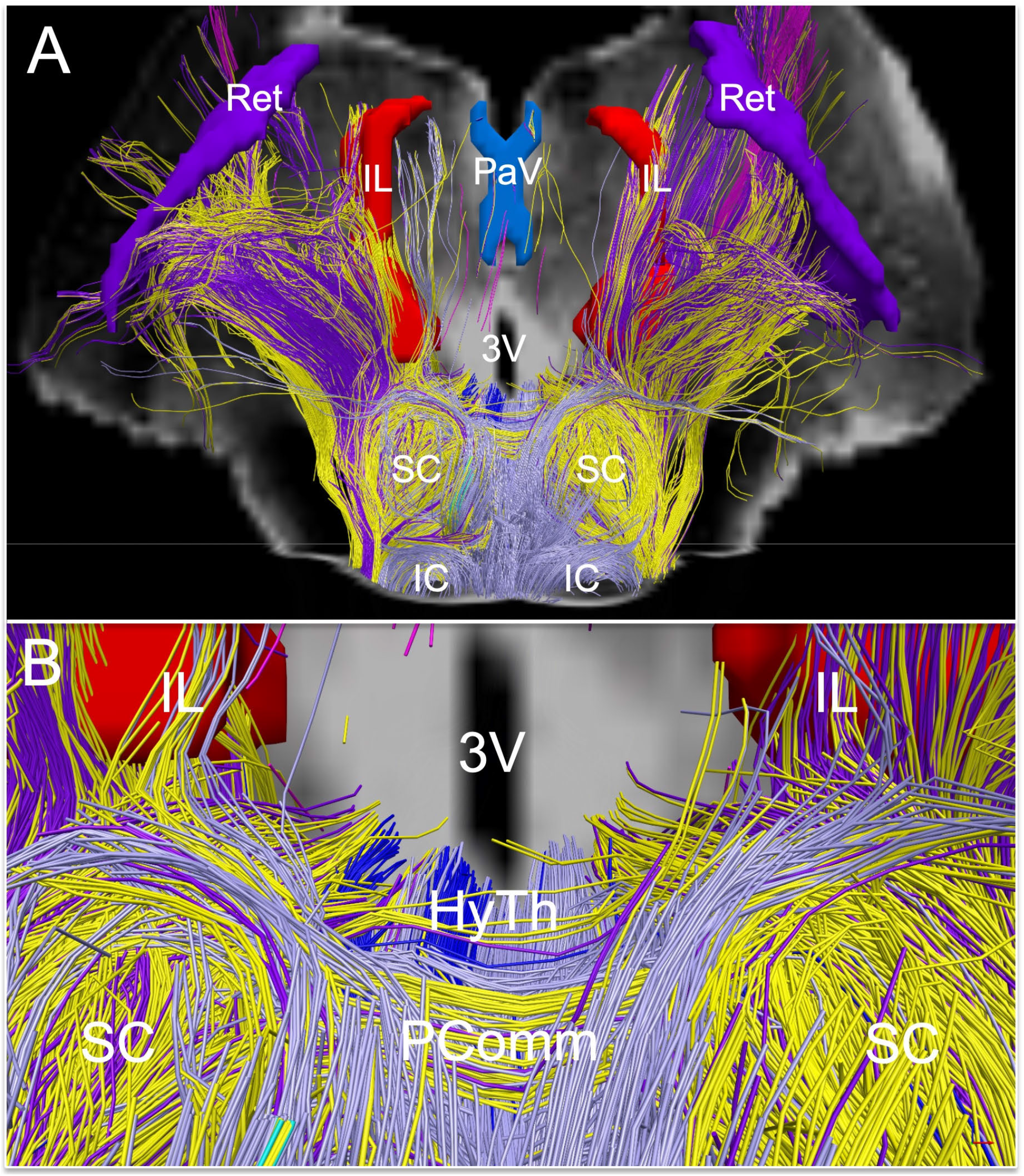
Extrareticular commissural and projection pathways of the default ascending arousal network. Deterministic fiber tracts emanating from extrareticular arousal nuclei are shown from a dorsal perspective in **(A)** and a zoomed dorsal perspective in **(B)**, superimposed on a coronal non-diffusion-weighted (*b=0*) image at the level of the mid-thalamus. Tract color-coding is identical to that in Fig. 3. Projection fibers are seen connecting with the hypothalamus (HyTh; individual hypothalamic nuclei not shown for clarity), and thalamic arousal nuclei: intralaminar nuclei (IL), paraventricular nuclei (PaV), and reticular nuclei (Ret). Commissural fibers cross the midline via the posterior commissure, with a high contribution of commissural fibers from the parabrachial complex (yellow), pedunculotegmental nucleus (dark purple), and periaqueductal grey (light purple).

### Functional connectivity between the dAAN and DMN

We used 7 Tesla resting-state fMRI (rs-fMRI) data from 84 healthy control subjects scanned in the Human Connectome Project (HCP) (*56*) to investigate the functional connectivity of the DMN with subcortical dAAN nodes. To map the DMN functional connectivity profile for each dAAN node, we used version 2.0 of the Harvard ascending arousal network (AAN) atlas (*9*) for brainstem nuclei. We used a previously published probabilistic segmentation atlas (*57*) for thalamic nodes, and an atlas proposed by Neudorfer et al. (*58*) for basal forebrain and hypothalamus nodes.

For each candidate brainstem, hypothalamic, thalamic, and basal forebrain node of the dAAN, we plotted the distributions of the DMN connectivity, as shown in Figure 8A. These analyses revealed a broad range of DMN connectivity within each candidate dAAN node. Several dAAN nodes contained voxels with positive correlations and negative correlations (i.e., anti-correlations) with the DMN. Within the brainstem, the VTA demonstrated the highest level of maximal DMN connectivity, while MnR showed the highest median level of DMN connectivity. Within the hypothalamus, thalamus and basal forebrain, the IL nucleus in the central thalamus demonstrated the highest level of maximal DMN connectivity, while the TMN nucleus in the hypothalamus showed the highest median level of DMN connectivity.

**Fig. 8.**
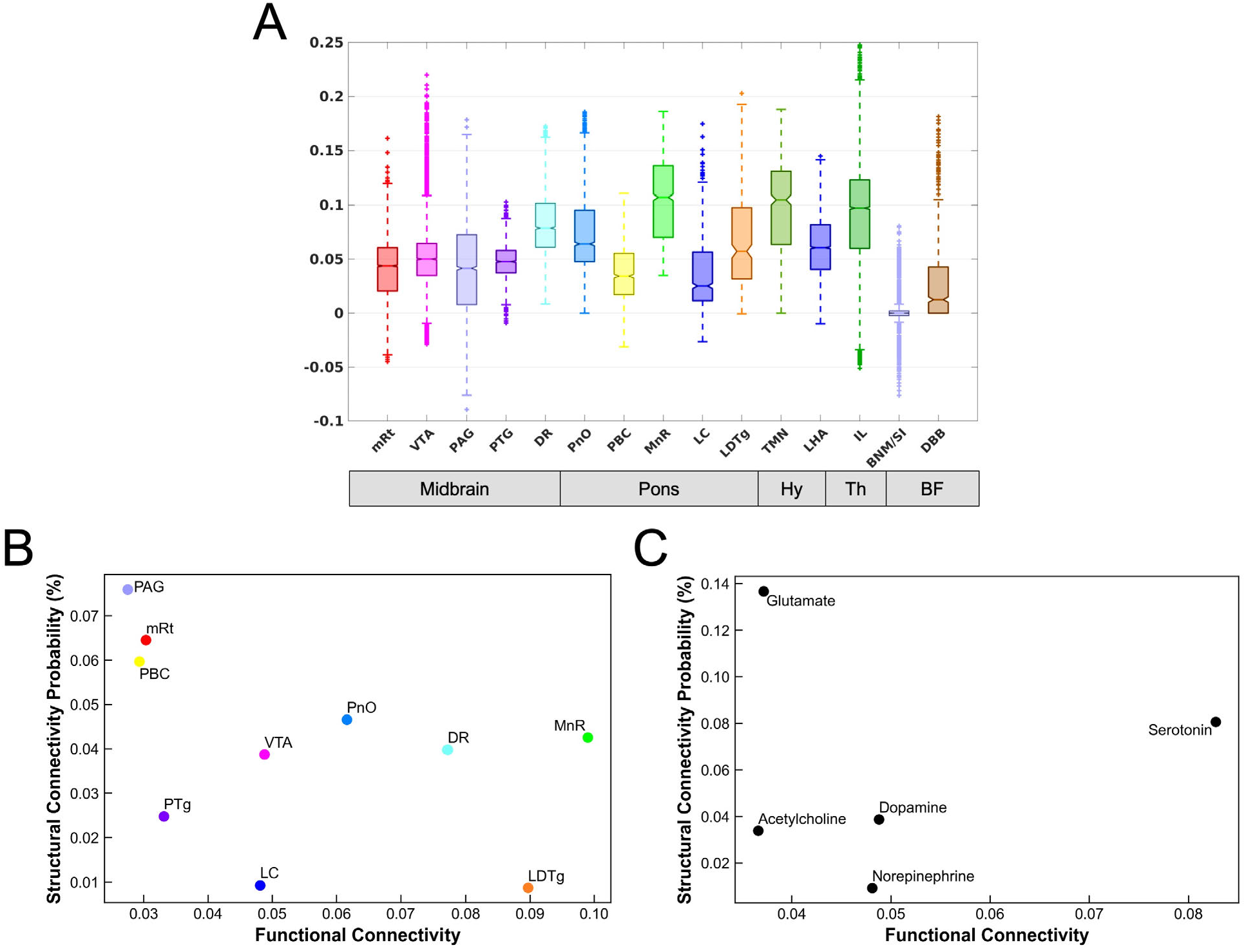
Functional connectivity between subcortical nodes of the default ascending arousal network and cortical nodes of the default mode network and their relationship to structural connectivity. (A) Boxplots of the DMN functional connectivity for each dAAN node. Y-axis represents the strength of functional connectivity (unitless) relative to the cortical DMN connectivity measured by the NASCAR method. VTA voxels showed the highest level of maximal functional connectivity with the DMN, while MnR had the highest level of median connectivity with the DMN. (**B**) Scatter plot between the mean of functional connectivity (x-axis) and the structural connectivity probability (y-axis) for each brainstem nucleus. (**C**) The counterpart grouped by the predominant type of neurotransmitter for each node: cholinergic = PTg and LDTg; dopaminergic = VTA; glutamatergic = PBC, mRt, and PnO; noradrenergic = LC; serotonergic = MnR and DR.

Given the complex relationship between structural and functional connectivity in cortical networks (59, 60), and the lack of data regarding structural-functional connectivity associations in subcortical networks, we plotted the relationship between functional connectivity mapped with in vivo 7 Tesla fMRI and structural connectivity mapped with ex vivo diffusion MRI (Figure 8B, 1. C) for connections between brainstem dAAN nodes and the cortical DM. In these analyses, the DMN was analyzed as a single target ROI. These structural-functional correlation plots revealed that several dAAN nodes, such as LDTg, MnR, and DR, had a high ratio of functional to structural connectivity, whereas other dAAN nodes, such as PAG, mRt, and PBC, had a high ratio of structural to functional connectivity. When grouping dAAN nodes by neurotransmitter system, serotonin and norepinephrine had high ratios of functional to structural connectivity, whereas glutamate had a high ratio of structural to functional connectivity.

## DISCUSSION

In this multimodal brain imaging study, we map the connections of a proposed subcortical network that sustains resting wakefulness in human consciousness. We provide evidence for a human default ascending arousal network (dAAN) comprised of interconnected nodes linking brainstem, hypothalamic, thalamic, and basal forebrain nuclei that are known from prior animal and human studies to sustain resting wakefulness. Further, we identified patterns of extensive structural and functional interconnectivity between subcortical dAAN nodes and cortical DMN nodes, providing a putative architecture for the integration of arousal and awareness in human consciousness. Consistent with prior human (*61*) and animal (*16, 18, 19, 62*) studies, convergent structural connectivity data from *ex vivo* diffusion MRI tractography and functional connectivity data from *in vivo* 7 Tesla resting-state fMRI revealed that a midbrain dAAN node – the dopaminergic VTA – is a widely connected hub node at the nexus of subcortical arousal and cortical awareness networks. Yet connectivity maps also revealed variable relationships between structural and functional connectivity of dAAN nodes, with serotonergic and noradrenergic nodes showing higher levels of functional connectivity with the DMN than would be expected based on their structural connectivity. These observations provide a basis for further investigation of the structural and functional brainstem properties that modulate the human DMN. We release all imaging data, immunohistochemistry data and network-based autopsy methods, including an updated atlas of dAAN brainstem nodes, to support efforts to map the connectivity of human consciousness.

### The VTA is a widely connected dAAN node

The VTA displayed extensive connectivity with other subcortical dAAN nodes and with cortical DMN nodes via multiple projection, association, and commissural pathways. Historically, the VTA was believed to mainly modulate behavior and cognition (*62*), but recent evidence from pharmacologic (*63, 64*), electrophysiologic (*16*), optogenetic (*18, 19*), and chemogenetic (*65*) methods, as well as behavioral experiments in dopamine knock-out mice (*66*), indicate that the VTA also modulates wakefulness. Functional connectivity studies from humans with disorders of consciousness further support the idea that the VTA modulates states of arousal, specifically via interactions with cortical DMN nodes (*17*). Our *ex vivo* diffusion MRI findings provide a structural basis for these functional observations in animals and humans, and our *in vivo* rs-fMRI findings provide further evidence for VTA-DMN connectivity, as VTA voxels showed the highest level of maximal functional connectivity of any brainstem dAAN node. These synergistic structural-functional connectivity findings solidify the role of dopaminergic VTA neurons in modulation of wakefulness, and hence consciousness (*67*).

Our observations supporting the role of the VTA as a potential dAAN hub node also reinforce ongoing efforts to target the dopaminergic VTA in therapeutic trials for patients with disorders of consciousness. Dopamine agonists and reuptake inhibitors are amongst the most commonly used pharmacologic therapies for patients with disorders of consciousness (*68*), and there is growing interest in using dopaminergic neuroimaging biomarkers to select patients for targeted therapies in clinical trials (*69-71*). The dAAN connectivity map and neuroimaging tools released here can thus be used to inform the design of future therapeutic trials and to expand the therapeutic landscape for patients with disorders of consciousness.

### Interconnectivity of the subcortical dAAN and cortical DMN in human consciousness

Connections between brainstem arousal nuclei have been demonstrated using a variety of tract-tracing and electrophysiological techniques in multiple animal species (*53, 72-76*), but prior evidence of such interconnectivity is limited in the human brain (*54, 77*). Here, we provide structural and functional connectivity evidence for multiple dAAN association pathways (i.e., connections between ipsilateral brainstem nuclei) and commissural pathways (i.e., connections between contralateral brainstem nuclei) that to our knowledge have not been previously visualized in the human brain. Moreover, we expand upon prior observations pertaining to projection pathways of human brainstem arousal nuclei (i.e., connections between brainstem nuclei and the thalamus, hypothalamus, basal forebrain, and cerebral cortex) (*9, 78-80*).

We identified four projection pathways that connect the human brainstem to the diencephalon and forebrain: the ventral tegmental tract, caudal division (VTT_C_); ventral tegmental tract, rostral division (VTT_R_); dorsal tegmental tract, medial division (DTT_M_); and the dorsal tegmental tract, lateral division (DTT_L_). The anatomic trajectories and connectivity patterns of these dAAN projection pathways are consistent with prior studies of human brainstem connectivity (*9, 54, 81, 82*) and are similar to their rodent (*53, 83*), feline (*76, 84*), and primate (*75*) homologues.

Our observation of direct connections between brainstem arousal nuclei and the cerebral cortex in the human brain are also consistent with extensive evidence in animal species (*74, 75, 85-90*). We found that the MFB and LFB are the primary pathways by which the human brainstem and cerebral cortex are connected, building upon prior observations about the human MFB (*54, 78, 91, 92*) and LFB (*54*). The human MFB travels through the hypothalamus and basal forebrain, carrying short-range tracts that connect with hypothalamic and basal forebrain nuclei via the VTT_C_. Long-range MFB fibers continue on to the cerebral cortex, connecting with DMN nodes within the frontal lobes. Although elucidation of the functional subspecialization of MFB and VTT_C_ pathways in the human brain awaits future investigation, these structural connectivity results are consistent with wake-promoting brainstem-hypothalamus-basal forebrain circuits in animals (*10*).

We also observed tracts diverging from the MFB within the hypothalamus and basal forebrain, then entering the anterior limb of the internal capsule and appearing to connect with the frontal cortex. Though these capsular tracts have been observed in prior human tractography studies (*78, 93*), their anatomic accuracy has been called into question by primate tract-tracing and immunostaining studies that show MFB capsular axons terminating in the basal ganglia, not the frontal cortex (*94*). Given the possibility that these capsular tracts connecting with the frontal cortex are “false positives” (i.e., anatomically implausible tracts), we do not assign them as being part of the MFB. Rather, all MFB connectivity results (Table 2) and anatomic labels (Figure 4) reported here are based on tracts that pass through the hypothalamus and basal forebrain before connecting with frontal DMN nodes.

We found that the LFB is the primary pathway by which human dAAN nodes connect with the temporo-parietal DMN nodes, including the PMC (i.e., posterior cingulate and precuneus), a hub node within the DMN whose connectivity is believed to be essential for conscious awareness (*11*). Functional connectivity between the VTA and the PMC was recently shown to modulate consciousness in patients with severe brain injuries and healthy volunteers receiving propofol sedation (*61*), observations that are supported by our VTA-PMC connectivity findings. Yet we also identified extensive DMN connectivity between multiple brainstem dAAN nodes beyond the VTA, with particularly strong connectivity in the PMC and MPFC nodes. Interestingly, midbrain dAAN nuclei showed more extensive connectivity with DMN nodes than did pontine dAAN nuclei. While a distance effect may partially account for this observation (i.e., the midbrain nodes are located in closer anatomic proximity to the DMN nodes), it is notable that three pontine nodes – PnO, PBC, and MnR – showed the highest connectivity with the MT node of the DMN in specimen 3. The functional significance of differential DMN connectivity patterns between midbrain and pontine dAAN nodes is unknown and requires future studies to evaluate the relative contributions of brainstem dAAN nuclei to modulating the DMN in the resting, conscious human brain.

### Anatomic location and neurotransmitter profiles of dAAN nodes

The 10 candidate brainstem dAAN nuclei that we identified from prior animal and human studies of arousal are all localized to the rostral pons and midbrain. The relationship between consciousness and brainstem nuclei in the rostral to mid-pons was demonstrated in Bremer’s classical *cerveau isole* experiments in cats (*95*) and subsequently confirmed in brainstem transection studies by Moruzzi (*96*), Steriade (*97*) and others. Similarly, lesions in the mid-pons, rostral pons, and midbrain have been observed to cause coma in humans (*98-102*), whereas lesions in the caudal pons and medulla do not. Although there is evidence for medullary modulation of arousal in animals (*103, 104*) and medullary connectivity with rostral brainstem arousal nuclei in humans (*54*), we did not find a definitive brainstem arousal nucleus caudal to the mid-pons that satisfied the *a priori* criteria for consideration as a candidate dAAN node. The analysis of medullary contributions to wakefulness and consciousness is beyond the scope of this paper and the subject of ongoing work.

Multiple candidate dAAN nodes contain GABAergic neurons, which modulate wakefulness via a variety of electrophysiological mechanisms. Amongst the glutamatergic neurons of the mesencephalic and pontine reticular formations (mRt and PnO), GABAergic neurons exert local inhibitory control upon the larger population of ascending stimulatory neurons (*105*). These GABAergic neurons may be responsible for the wake-promoting behavioral effects of PnO via local inhibition (i.e., “gating”) of REM sleep-promoting cholinergic neurons (*38*). Moreover, GABAergic neurons of the thalamic reticular nucleus (Ret) transition from intermittent burst-firing to tonic firing during the transition from slow-wave sleep to wakefulness (*106*). We found extensive connectivity between Ret and other dAAN nodes, consistent with prior electrophysiologic studies showing that Ret neurons synapse widely with ascending brainstem arousal inputs and receive inputs from arousal neurons of the basal forebrain (*107, 108*). Our connectivity findings thus reinforce the key role of GABAergic modulation of arousal in the human brain.

### Applications of dAAN connectivity maps for patients with disorders of consciousness

We expect that the network-based autopsy methods and connectivity data generated here will advance the study of coma pathogenesis and recovery (*1*). Classification of projection, association, and commissural pathways within the human dAAN will allow categorization of different combinations of preserved pathways in patients who recover from coma. This effort may ultimately enable identification of the minimum ensemble of nodes and connections that are sufficient for sustaining wakefulness in human consciousness. In addition, the dAAN connectivity measures have the potential to inform prognostication and patient selection for therapeutic trials aimed at stimulating the dAAN and promoting recovery of consciousness (*69*).

Reemergence of consciousness after severe brain injury does not require the entire set of grey matter nodes and white matter connections that sustain wakefulness in the healthy human brain. Detailed clinico-anatomic studies of patients with focal brainstem lesions and brainstem axonal disconnections who recovered wakefulness after a period of coma convincingly illustrate this point (*82, 98, 102, 109, 110*), as do a multitude of animal studies showing that recovery of wakefulness is possible after experimental lesioning of brainstem or diencephalic arousal nuclei(*111*). Even in the classical *cerveau isole* experiments, in which cats are initially comatose due to transverse transection of the brainstem at the intercollicular level of the midbrain (*112*), some cats ultimately recover behavioral and electrophysiological evidence of wakefulness after several weeks (*113*). Similarly, prolonged coma (i.e., ≥ 4 weeks) in human survivors of severe brain injury is extremely rare (*114*). Thus, identification of ensembles of dAAN connections that are compatible with recovery and receptive to therapeutic stimulation has potential to improve outcomes for patients with severe brain injuries (*115*).

The precise mechanisms by which residual connections enable reemergence of wakefulness after coma are unknown, but observational and therapeutic studies are beginning to identify specific subcortical circuits that mediate recovery of wakefulness after severe brain injury (*1, 82, 116*). Complementary evidence from neurophysiologic and cell culture studies suggests that neuronal networks can adapt to perturbations and reestablish normal function (*117-119*). These plasticity mechanisms provide a basis for functional reorganization of subsets of dAAN nodes in response to structural injury, suggesting that consciousness can be reestablished by variable combinations of network components (*1*).

These potential mechanisms of dAAN plasticity in humans recovering from coma are also consistent with theoretical models of the resilience and adaptability of neural systems. Modeling experiments suggest that neural systems with high complexity have high measures of degeneracy and redundancy, meaning that different elements within the system can perform the same function (*120*). These model-based observations are supported by human therapeutic studies in patients with disorders of consciousness (*115*), which suggest that there are multiple targets for pharmacologic or electrophysiologic reactivation of arousal circuits (*17, 23, 121-125*). Considering animal evidence for electrical coupling of arousal nuclei (*126*), one interpretation of these human therapeutic trials is that stimulation of different network nodes may lead to a final common pathway of increased functional coupling within the entire arousal network.

### Limitations and methodological considerations

In deterministic and probabilistic tractography, a fiber tract represents an inferential model of axonal pathways based upon measurements of directional water diffusion along these pathways. The number of axons that corresponds to a tract depends partly on the spatial resolution (i.e., voxel size) of the diffusion data. In the experiments reported here, the voxel size was almost 1,000 times larger in dimension than the diameter of a single axon (∼600-750 microns versus ∼1 micron). Hence, the tractography results reported here do not represent “ground truth” axonal anatomy. All tractography results should be considered inferential and are subject to the inherent methodological limitations of diffusion MRI tractography, which have been well characterized (*127-131*).

Given these methodological limitations, our tractography-based definition of anatomic connectivity does not prove synaptic connectivity, which will require confirmation in future studies that move from the mesoscale connectome (i.e., ∼600 to 750 µm resolution) to the microscale synaptome (i.e., ∼1-10 µm), a major area of ongoing study in neuroscience (*132*). However, there are currently no available methods to integrate our mesoscale human connectome map with prior microscale animal synaptome wiring diagrams (*73*). Innovations such as CLARITY (*133*) and polarization-sensitive optical coherence tomography (*134*) provide hope for this endeavor, but an integrated connectome-synaptome map of the dAAN is currently out of reach.

Another limitation is that diffusion tractography cannot determine the direction of electrical signaling within a tract. There is extensive electrophysiological evidence for “top-down” modulation of arousal (*135, 136*) and cognition (*137*), and the MFB is known to carry descending fibers alongside its ascending fibers (*138*). In addition, multiple types of neurons within the arousal network may have bifurcating axons with ascending and descending projections (*89*). Thus, our use of the term “ascending” is not intended to suggest that arousal is mediated solely by ascending pathways from the brainstem to rostral sites. Rather, homeostatic functions such as arousal are modulated by bi-directional “bottom-up” and “top-down” signaling between the brainstem, diencephalon, basal forebrain, and cerebral cortex.

It is also important to consider that false positive tracts can be generated by a “highway-effect” whereby tracts that run alongside one another can merge into anatomically implausible pathways. Conversely, false negative findings can occur due a variety of factors, including increased distance between nodes and tractography’s low sensitivity for identifying axons with multidirectional branching patterns (*139, 140*), small diameters (*75*), and/or lack of myelination (*75*). These latter anatomic features are known characteristics of the rostral serotonergic, noradrenergic, and cholinergic axonal projections, which may explain the absent or uncertain connectivity findings for MnR, LC, and LDTg (see Table 2). We attempted to mitigate these problems by performing dMRI with a spatial resolution of 600-750 µm and an angular resolution of 60-90 diffusion-encoding directions. Further, we tested for connectivity between each candidate dAAN node and an anatomically implausible control node to empirically derive a “noise window”, and we only considered connections that exceeded this statistical threshold. Yet even with high-resolution data and strict statistical thresholds, the possibility of false positive and false negative structural connectivity remains.

We also acknowledge that the present study does not address functional subspecialization, or fractionation, of different circuits within the dAAN. Given that dAAN nodes contain neurons utilizing multiple neurotransmitters (*7*), there is substantial intra-nodal functional complexity that is not accounted for in our structural connectivity methods, which treat each node as a singular entity. For example, the PnO contains neurons involved in activation of REM sleep (*141*), generation of the hippocampal theta rhythm (*142*), and active movement during waking (*143*), but here we considered the PnO solely as an arousal nucleus because of its population of wake-promoting GABAergic neurons (*38*). Similarly, the PTg contains diverse populations of cholinergic, glutamatergic, and GABAergic neurons with different roles in modulation of wakefulness (*144*). Nodal parcellation of the PnO, PTg and other multi-function nodes will require future dAAN fractionation studies, just as fractionation of DMN nodes was performed in the years following the DMN’s discovery (*145, 146*).

### Conclusions and future directions

We provide insights into the connectivity of a subcortical arousal network that sustains resting wakefulness in the conscious human brain. We found that subcortical dAAN nodes interconnect with cortical DMN nodes, providing a neuroanatomic basis for integration of arousal and awareness in human consciousness. The dAAN connectivity map and neuroanatomic taxonomy proposed here aims to facilitate further study into the role of subcortical arousal pathways in human consciousness and its disorders. The small size and neuroanatomic complexity of the human brainstem renders it inherently difficult to study, but as new neuroimaging tools are developed to investigate human brainstem connectivity (*147-152*), we expect that it will be possible to refine this dAAN connectivity map and definitively identify the networks that sustain human consciousness.

## MATERIALS AND METHODS

### Identification of candidate dAAN nodes

We identified candidate dAAN nodes via four lines of previously published evidence: electrophysiological, gene expression, lesion, and stimulation. Specifically, we identified subcortical nuclei for which prior studies demonstrated any one of the following: 1) tonic activity during resting wakefulness (*28, 153*); 2) increased Fos staining after periods of arousal (*154*); 3) lesion-induced loss of resting wakefulness (*95*); or 4) stimulation-induced arousal (*13*) (see **Table 1**). Animal electrophysiology, gene expression, lesion, and stimulation studies were identified using reference lists from text books (*28, 111, 155-158*), review articles (*10, 106, 159-162*), and from the authors’ reference libraries.

These *a priori* criteria do not stipulate that dAAN nodes must release excitatory neurotransmitters. Although the overall influence of the dAAN on the cerebral cortex is activating, multiple lines of theoretical and experimental evidence indicate that inhibitory circuits are essential for the modulation and stability of activating networks (*158, 163-165*). At a circuit level, GABAergic neurons exert multiple types of inhibition, including feedback, feed-forward and lateral inhibition. These inhibitory mechanisms enhance the nonlinearity and complexity of neural networks, and hence their resiliency and stability during perturbations (*28*). Of particular relevance to wakefulness, *in vivo* and *in vitro* animal studies suggest that GABAergic neurons coordinate the synchronous fast activity of subcortical arousal neurons (*7, 135, 162*). We therefore consider nodes containing GABAergic inhibitory neurons as candidate dAAN nodes.

Of note, the nodes and connections of the dAAN need not be the same as those that facilitate state changes. For example, the hypothalamic and brainstem nodes of the subcortical “flip-flop switch” that mediates transitions between sleep and wake (*161*) are not necessarily the same nodes that sustain wakefulness once it is established. Rather, the flip-flop switch may activate a distinct or overlapping set of dAAN nodes once the transition from sleep to wakefulness has occurred. Here, we focus exclusively on the nodes and connections of the dAAN that sustains the resting mode of the wakeful state.

### Brain specimen acquisition and processing

Written informed consent for brain donation was obtained postmortem from family surrogate decision–makers, in accordance with a research protocol approved by the Institutional Review Board at Mass General Brigham. Criteria for brain specimens being included in the study were: 1) no history of neurological illness; 2) no abnormal *in vivo* brain scan; 3) normal neurological examination documented by a clinician in the medical record prior to death; and 4) normal postmortem gross examination of the brain by a neuropathologist. All specimens were fixed in 10% formaldehyde.

### Ex vivo MRI data acquisition

For Specimen 1, we dissected the hemispheres from the diencephalon and basal forebrain, and dissected the cerebellum from the brainstem to fit the specimen into the small bore of a 4.7 Tesla MRI scanner, as previously described (*9*). The dissected specimen consisted of the rostral pons, midbrain, hypothalamus, thalamus, basal forebrain, and part of the basal ganglia. Specimens 2 and 3 were scanned as whole brains on a large-bore 3 Tesla MRI scanner. Immediately prior to scanning, the specimens were transferred from 10% formaldehyde to Fomblin (perfluropolyether, Ausimont USA Inc., Thorofare, NJ), which reduces magnetic susceptibility artifacts (*166*).

#### Diffusion MRI of dissected specimen

Specimen 1 was scanned in a small horizontal-bore 4.7 Tesla MRI scanner (Bruker Biospec), as previously described (*9*). We utilized a 3-dimensional diffusion-weighted spin-echo echo-planar imaging (DW-SE-EPI) sequence, with 60 different diffusion-weighted measurements at *b* = 4057 s/mm^2^. One dataset of 3D DW-SE-EPI data with *b* = 0 s/mm^2^ was also acquired to calculate quantitative diffusion properties in each voxel. Additional sequence parameters were: gradient strength = 12.4 G/cm, duration ∂ = 13.4 msec, intertemporal pulse offset Δ = 25 msec, repetition time (TR) = 1000 msec, echo time (TE) = 72.5 msec, and spatial resolution = 562 x 609 x 641 µm. Total diffusion scan time was 2 hours and 10 minutes.

#### Diffusion MRI of whole brain specimens

Specimens 2 and 3 were scanned as whole brains on a 3 Tesla Tim Trio MRI scanner (Siemens Medical Solutions, Erlangen, Germany) using a 32-channel head coil. To increase diffusion sensitivity on the large-bore 3 Tesla MRI clinical scanner that was required to scan a whole brain specimen, diffusion data were acquired using a 3D diffusion-weighted steady-state free-precession (DW-SSFP) sequence (*47, 167*). DW-SSFP is an SNR efficient approach that can achieve diffusion weighting without requiring long TE, making it well suited for *ex vivo* samples characterized by short T2 (*168*). We acquired 90 diffusion-weighted volumes and 12 b=0 volumes at 750 µm isotropic spatial resolution. Total diffusion scan time was 30 hours and 31 minutes per specimen. In contrast to the classic DW-SE-EPI sequence utilized for specimen 1, the diffusion-weighting for a DW-SSFP sequence is not defined by a single global *b* value, as it depends on other tissue and imaging properties, such as the T1 relaxation time, the T2 relaxation time, the TR, and the flip angle (*167*). Using the formulation in (*168*) and previously reported *ex vivo* estimates of T1, T2, and diffusivity (D) (T1= 400 ms, T2=45 ms, D=0.08×10^-3^ mm^2^/s) we computed an “effective” b-value of 3773 s/mm^2^ (*169*).

### Cortical surface parcellation of whole brain specimens

We parcellated the cortical nodes of the DMN in whole brain specimens 2 and 3 for the dAAN-DMN connectivity analysis. DMN nodes were parcellated according to the canonical Yeo 7-network atlas (*170*), as instantiated in the 1000-node Schaefer version of the atlas (*171*) available at https://github.com/ThomasYeoLab/CBIG/tree/master/stable_projects/brain_parcellation/Schaefer2018_LocalGlobal/Parcellations/project_to_individual.

First, to generate cortical surfaces for identification of DMN nodes, we acquired multi-echo flash (MEF) data at 1 mm isotropic resolution on the 3 Tesla Tim Trio MRI scanner during the same scanning session during which the diffusion data were acquired. Key MEF sequence parameters were TR = 23.00 msec, TE = 2.64 msec, and 6 flip angles of 5° to 30° were collected at 5° increments. Synthetic T1 and proton density (PD) maps were estimated directly from the MEF acquisitions using the DESPOT1 algorithm (*172*), available through the FreeSurfer (http://surfer.nmr.mgh. harvard.edu) utility “mri_ms_fitparms” (*173*).

#### Cortical surface generation and neuroanatomic localization of DMN nodes

To generate the cortical surface model, the synthesized PD volume was processed to remove extraneous background noise by thresholding the PD volume to create a mask containing hyperintense voxels. The mask was then subtracted from the initial PD volume to achieve a more homogenous image. Using the intensity-corrected PD volume, we then generated white and grey matter segmentations using the modified Sequence Adaptive Multimodal Segmentation (SAMSEG) pipeline (*174*). The SAMSEG white and grey matter segmentations were then used to generate the pial and white matter surfaces.

Once the surfaces were generated, we performed surface-based atlas registrations to create labels corresponding to the DMN nodes in the canonical Yeo atlas: posteromedial complex (PMC; i.e., posterior cingulate cortex and precuneus), medial prefrontal cortex (MPFC), inferior parietal lobule (IPL), lateral temporal lobe (LT), and medial temporal lobe (MT). To translate the surface-based MEF labels into volumetric diffusion space we used the boundary-based registration (*175*) method available through FreeSurfer. Using the transformation matrix obtained through coregistration, we then translated our atlas labels from MEF surface space to volumetric diffusion space for further analysis. We then projected the Yeo atlas labels onto those surfaces using a fixed 2 mm distance into the surface via a nonlinear spherical transform (*176*).

Detailed processing parameters are provided in our code, which we release at https://github.com/ComaRecoveryLab/Network-based_Autopsy. All data were processed using a standard release of FreeSurfer 7.1.2 available at https://github.com/freesurfer/freesurfer and FSL6.0.3 available at https://fsl.fmrib.ox.ac.uk/fsl/fslwiki.

### Sectioning and staining for histologic-radiologic correlations

Upon completion of *ex vivo* MRI data acquisition, specimen 1 was divided into seven blocks, which were embedded in paraffin. We then cut serial transverse sections of the rostral pons and midbrain, and coronal sections of the hypothalamus, thalamus, and basal forebrain at 10 µm thickness (LEICA RM2255 microtome, Leica Microsystems, Buffalo Grove, IL, USA). Every 50^th^ section was stained with hematoxylin-and-eosin for nuclei and counterstained with Luxol fast blue for myelin, yielding a stained section every 500 µm. The purpose of these stains was to define the neuroanatomic location, borders, and morphology of all candidate dAAN nodes.

Additionally, 10 µm sections from the midbrain were prepared using tyrosine hydroxylase (Pel-Freez Biologicals; rabbit polyclonal P40101) counterstained with hematoxylin, and 10 µm sections from the pons were prepared using tryptophan hydroxylase (Sigma-Aldrich; mouse monoclonal AMAB91108) counterstained with hematoxylin. Tyrosine hydroxylase more precisely characterizes the location, borders and morphology of the dopaminergic ventral tegmental area (VTA), as detailed in the **Supplementary Material**. Tryptophan hydroxylase characterized the location, borders and morphology of the serotonergic median raphe (MnR) and dorsal raphe (DR).

### Histology-guided neuroanatomic localization of brainstem dAAN nodes

For specimen 1, candidate dAAN nodes (i.e., nodes selected based on the *a priori* criteria) were traced on the diffusion-weighted images using histological guidance. Each diffusion image was compared to its corresponding histological section to ensure that radiologic dAAN nodes shared the same location and morphology as the histological dAAN nuclei (**Figs. S4, S5**). Histological nuclei were delineated by visual inspection of their cytoarchitecture with a light microscope and with digitized slides. We then cross-referenced with the Paxinos atlas of human brainstem neuroanatomy for brainstem dAAN nodes (*51*) and with the Allen Institute atlas of the human brain for thalamic, hypothalamic, and basal forebrain dAAN nodes (*52*). Since brainstem nuclei change shape, size and contour along the rostro-caudal axis, histological guidance of dAAN node tracing was performed for every axial diffusion image using its corresponding histological section, yielding a 3-dimensional model of brainstem arousal nuclei.

For specimens 2 and 3 the brainstem dAAN nuclei were also traced manually on each diffusion MRI dataset, but with reference to canonical atlases and selected immunostains, not serial immunostains or comprehensive histological data. Specifically, the neuroanatomic boundaries of brainstem nodes were visually confirmed with respect to two reference templates: 1) the *ex vivo* template of histologically guided nodes generated for specimen 1, which we release here as part of the Harvard Disorders of Consciousness Histopathology Collection (http://histopath.nmr.mgh.harvard.edu); and 2) the Paxinos human brainstem atlas (*51*).

### Histology-guided neuroanatomic localization of diencephalic and forebrain dAAN nodes

Similar to the brainstem nucleus segmentation, we used histological guidance via coronal sections through the diencephalon and basal forebrain to identify the borders of thalamic, hypothalamic, and basal forebrain dAAN nodes on the diffusion MRI dataset for Specimen 1. To segment the thalamic, hypothalamic, and basal forebrain nuclei for specimens 2 and 3, we manually traced each node based on the Allen Institute human brain atlas (*52*).

### Overview of *ex vivo* diffusion MRI diffusion data processing for tractography

We processed the *ex vivo* diffusion datasets for deterministic and probabilistic tractography analyses to generate insights into both qualitative anatomic relationships of dAAN tracts and quantitative dAAN connectivity properties. Deterministic tractography yields more visually informative qualitative data about the neuroanatomic relationships between tracts (e.g., sites of tract branching and crossing) (*54, 151*), whereas probabilistic tractography provides a more reliable quantitative measure of connectivity between nodes because it accounts for uncertainty in tract trajectories and can detect tracts on and past grey matter boundaries (*177*).

Diffusion data from all three specimens were processed for deterministic fiber tracking, and diffusion data from specimens 2 and 3 were processed for probabilistic fiber tracking. Specimen 1 was excluded from the probabilistic tractography analysis because it was scanned using different acquisition parameters, which precludes comparison of quantitative probabilistic data with Specimens 2 and 3 without complex modelling that falls outside of the scope of these experiments. To optimize consistency across methods, we standardized the deterministic and probabilistic processing parameters wherever possible. For example, we did not apply an FA threshold to the deterministic or probabilistic analyses because low FA has been observed in the rostral brainstem tegmentum (*9, 178*), and thalamus (*179*) in the human brain. We confirmed this observation in an FA analysis of tracts emanating from the mRt in specimen 1, which showed low FA in multiple segments of mRt tracts passing through the brainstem tegmentum (**Fig. 2**. These low FA measurements may be attributable to the complex meshwork of crossing fibers within the brainstem tegmentum (*84*), varying degrees of myelination within brainstem axons (*75*), and/or nuclei that are interspersed amongst the axons (*51*). Additional details regarding deterministic and probabilistic processing parameters, such as the selection of angle thresholds and the rationale for not applying a distance correction, are provided in the **Supplementary Material**.

### Deterministic tractography analysis of dAAN connectivity

For deterministic tractography, diffusion data for all three specimens were processed in Diffusion Toolkit 6.3 (*180*). Orientation distribution functions (ODFs) for each voxel were obtained utilizing a spherical harmonic q-ball reconstruction approach (*181*) and whole brain tractograms were generated using the FACT algorithm and the ODF’s principle diffusion directions (*182*). Q-ball reconstruction accounts for the possibility of multiple diffusion directions within each voxel, thus enhancing detection of crossing, splaying, and/or “kissing” fibers within complex neuronal networks (*183, 184*). DAAN tracts were isolated manually in TrackVis, version 6.1 (*180*), using each candidate dAAN nucleus as an inclusion region of interest. Tracts were visually analyzed to determine the trajectories, branching patterns, and anatomic relationships between bundles connecting the candidate dAAN nodes as well as dAAN-DMN connections.

### Probabilistic tractography analysis of dAAN connectivity

For probabilistic tractography, diffusion data for the two whole-brain specimens (specimens 2 and 3) were processed using FMRIB’s Diffusion Toolbox, version 6.0.6. Estimates of principle diffusion orientation in each voxel were obtained using the ball-and-stick model (*177, 185*) in bedpostx with two stick compartments. Tractography was performed using probtrackx with default parameters. Connectivity was analyzed between all candidate dAAN nodes using the “network” flag. For each pair of nodes, 5,000 tracts (i.e., streamlines) were propagated from each voxel within the seed node, and the target node was treated as both a “termination mask” and a “waypoint mask” (*54, 151*).

### Statistical Analysis of Connectivity

We computed an undirected connectivity probability (*CP*) for each pair of nodes (i.e., node A and node B) by performing two analyses: 1) node A was treated as a seed and node B as a target; and 2) node B was treated as a seed and node A as a target. Using this symmetric connectivity model (*186*), we computed *CP_A→B_*, *CP_B→A_*, and *CP_A↔B_*:

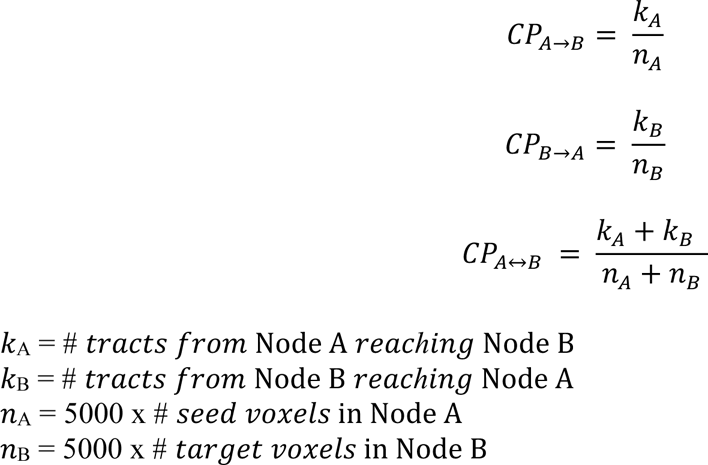

*CP_A↔B_* hence represents the probability of a tract connecting node A and node B, weighted by the number of voxels within each node and invariant to direction (*81, 82*). Importantly, the symmetric model reduces a potential statistical bias in probabilistic tractography: the probability of a connection between node A and node B being underestimated if node A has widely distributed connectivity. The probability of a tract connecting with node B would therefore decrease as the probability of that tract connecting with other nodes increases. The symmetric model mitigates, though does not eliminate, this potential bias.

Although complete elimination of false-positive tracts (i.e., anatomically implausible connections) is not possible with either probabilistic or deterministic tractography, we attempted to account for the possibility of false-positive tracts by comparing *CP* values for each candidate brainstem dAAN node with two control nodes: the basis pontis (BP) and the red nucleus (RN). Based on prior anatomic labeling studies in animal models, BP and RN are not expected to have significant connectivity with the candidate brainstem dAAN nodes (e.g., (*53*)), as they mainly contain cortico-ponto-cerebellar fibers that connect the cerebellum to the primary motor cortex (and vice versa), the primary function of which is motor activity (*187*). We established two *a priori* criteria, both of which must be met, for operationally defining a structural connection between a pair of candidate dAAN nodes (nodes A and B): 1) *CP_A↔B_* > 0; and 2) the 95% confidence interval (CI) for *CP_A↔B_* must not overlap with the 95% CI for any of the following-*CP_A↔BP_, CP_A↔RN_*, *CP_B↔BP_*, and *CP_B↔RN_*.

The 95% CI for each *CP* measurement was calculated by modeling *CP* as a binomial distribution:

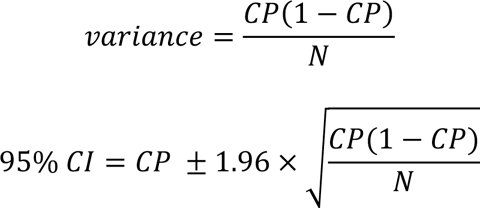

where *N* = *n_A_* + *n_B_*.

The binomial statistical model depends upon the assumption that the success of each trial (i.e., the likelihood that a tract from the seed will connect with the target) is independent from prior trials. This assumption may not always be valid, because there is a possibility of intra-voxel dependence (i.e., the success of tract 1 launched from voxel 1 may be dependent on the success of tract 2 launched from voxel 1) and inter-voxel dependence (i.e., the success of tract 1 launched from voxel 1 may be dependent on the success of tract 1 launched from voxel 2). Nevertheless, deriving a quantitative model of these dependencies would require accounting for multiple anatomic factors that likely vary from node to node, or even from voxel to voxel, and is therefore beyond the scope of the present study. Crucially, the binomial model likely overestimates variance because it does not account for the aforementioned dependencies. Hence, we believe that the binomial model implemented here is a conservative approach for estimating *CP* variance and the 95% CI.

### Proposed neuroanatomic taxonomy for the human dAAN

We classified dAAN connections as projection, association, or commissural. This classification system is intended for use as a neuroanatomic taxonomy for the subcortical dAAN, akin to that of cortical networks whose connections are classified in these terms (*55*). In the cortical classification system, the cortex is the frame of reference for defining fiber pathways. Projection fibers project from the cortex to subcortical regions. Association fibers connect ipsilateral cortical regions, and commissural fibers cross the midline to connect contralateral cortical regions.

Here, we propose that the brainstem is the frame of reference for classifying fiber pathways of the dAAN. As such, projection fibers are defined as those that project from brainstem nuclei to rostral diencephalic nuclei, basal forebrain nuclei, and/or cortical regions that mediate consciousness (i.e., DMN nodes). Association fibers connect brainstem nuclei to ipsilateral brainstem nuclei, and commissural fibers cross the midline to connect with contralateral brainstem nuclei.

### 7 Tesla resting-state functional MRI of dAAN-DMN connectivity

7 Tesla resting-state fMRI (rs-fMRI) scans from 84 healthy controls in the Human Connectome Project (HCP) (*56*) were used in this study. The rs-fMRI data were minimally preprocessed, resampled, and co-registered to the MNI template by the HCP pipeline (*188*). Grayordinate representations of the data were used for joint analysis of cortical and subcortical connectivity, where cortical data were sampled onto the subject surface, and subcortical data were represented in volumetric space (*188*). We did not apply any further spatial smoothing to the minimally preprocessed data because typical Gaussian smoothing would mix signals across different functional regions (*189*), which may cause a significant issue when resolving relationships between nuclei in dAAN (*190*). Instead, the improved SNR was obtained using a tensor-based group analysis, as described below.

Using our previously developed pipeline (*191*), we estimated the functional connectivity from dAAN to the canonical DMN. Specifically, we first temporally synchronized the rs-fMRI data of each subject to a group average using the group BrainSync algorithm (*192, 193*). We further arrange the synchronized data from all subjects along the third dimension (the first dimension is space, and the second dimension is time), forming a 3D data tensor. We then applied the NASCAR tensor decomposition method (*194, 195*) to discover the whole-brain DMN. The subcortical section was separated from the grayordinate structure and converted to a volumetric NIFTI file.

To extract the connectivity profile for each nucleus in dAAN, we used version 2.0 of the Harvard Ascending Arousal Network (AAN) Atlas (*9*) for brainstem nuclei, the probabilistic segmentation atlas (*57*) for thalamic nuclei, and an atlas proposed by Neudorfer et al. (*58*) for basal forebrain and hypothalamus nuclei to define the regions of interest (ROI). We then, for each ROI, plot the distributions of the DMN connectivity values using box plots. Of note, three candidate diencephalic dAAN nodes – the supramammillary nucleus of the hypothalamus (SUM), the paraventricular nucleus of the thalamus (PaV), and the reticular nucleus of the thalamus (Ret), were not included in the MRI atlases.

## SUPPLEMENTARY MATERIALS

### Abbreviations

AAN: ascending arousal network
ARAS: ascending reticular activating system
CEM/Pf: centromedian/parafascicular nucleus of the thalamus
CHN: central homeostatic network
CL: central lateral nucleus of the thalamus
CnF: cuneiform nucleus
CTT: central tegmental tract
dAAN: default ascending arousal network
DBB: diagonal band of Broca
DMN: default mode network
DR: dorsal raphe
DTT: dorsal tegmental tract
DTT_L_: dorsal tegmental tract, lateral division
DTT_M_: dorsal tegmental tract, medial division
DWI: diffusion-weighted imaging
EEG: electroencephalography
FA: fractional anisotropy
fMRI: functional magnetic resonance imaging
IC: inferior colliculus
LC: locus coeruleus
LDTg: laterodorsal tegmental nucleus
LFB: lateral forebrain bundle
LHA: lateral hypothalamic area
MFB: medial forebrain bundle
ML: medial lemniscus
MLF: medial longitudinal fasciculus
MnR: median raphe
MPFC: medial prefrontal cortex
MRI: magnetic resonance imaging
mRt: mesencephalic reticular formation
NBM: nucleus basalis of Meynert
PAG; periaqueductal grey; PBC: parabrachial complex
PBP: parabrachial pigmented nucleus of the ventral tegemental area
PComm: posterior commissure
PIF: parainterfascicular nucleus
PMC: postero-medial complex (i.e. posterior cingulate and precuneus)
PN: paranigral nucleus of ventral tegmental area
PnO: pontine reticular nucleus = oral part (pontis oralis)
PTg: pedunculotegmental nucleus
PV: paraventricular nucleus of the thalamus
Ret: reticular nucleus of the thalamus
RIs: retroisthmic nucleus
RLi: rostral linear nucleus
RMg: raphe magnus
SC: superior colliculus
SCP: superior cerebellar peduncle
SI: substantia innominata
SN: substantia nigra
TM: tuberomammillary nucleus
VLPO: ventrolateral pre-optic area
VTA: ventral tegmental area
VTAR: ventral tegmental area, rostral part
VTT: ventral tegmental tract
VTT_C_: ventral tegmental tract, caudal division
VTT_R_: ventral tegmental tract, rostral division.

## SUPPLEMENTARY METHODS

### A representative animal electrophysiological study of arousal

In 1982, Steriade and colleagues tested the hypothesis generated by Moruzzi and Magoun in 1949 that the brainstem reticular formation drives tonic activation of thalamocortical circuits, and therefore cortical EEG desynchronization, by increasing firing rates during wakefulness versus slow-wave sleep (*35*). Rather than using electrical stimulation or lesion-based techniques, Steriade and colleagues recorded extracellularly the spontaneous firing of mRt neurons while seven cats were awake, sleeping, or transitioning between these states, as determined by cortical EEG patterns. MRt neuron firing rates were significantly higher during wakefulness than sleep. This finding was most prominent in mRt neurons projecting rostrally versus caudally (i.e., to the paramedian pons). Moreover, the increase in firing rate corresponded to EEG and behavioral evidence of wakefulness and was tonic in nature, suggesting a direct link between mRt activity, activation of the cortical EEG, and behavioral evidence of wakefulness. These results suggest that the mRt plays an important role in sustaining wakefulness as part of the dAAN.

### Overview of neuroanatomic localization of candidate dAAN nodes

To facilitate reproduction of our methods, we describe the neuroanatomic boundaries of the candidate brainstem dAAN nodes in **Tables S1-S5**. In addition, we detail our approach to defining the neuroanatomic borders of each nucleus. We use the Paxinos atlas as our primary reference (*48*), but we also consider differences between the Paxinos atlas and the Olszewski and Baxter brainstem atlas (*50*), another canonical brainstem atlas of human neuronanatomy. Where applicable, we explain our rationale for mediating discrepancies in anatomic borders and/or nomenclature between these atlases, as well as between these atlases and other neuroanatomic studies. Finally, we explain for each brainstem dAAN node how the nomenclature and borders proposed here differ from those that our laboratory has used previously (*9, 54, 99, 109, 151*), including in our prior analyses of Specimen 1 (*9, 99*). At the completion of this process, we updated the Harvard Ascending Arousal Network Atlas and now release version 2.0 of the atlas (doi:10.5061/dryad.zw3r228d2; download link: https://datadryad.org/stash/share/aJ713eXY12ND56bzOBejVG2jmOFCD2CKxdSJsYWEHkw).

### Neuroanatomic localization of midbrain dAAN nodes

#### Mesencephalic Reticular Formation (mRt)

The mRt is a region of the midbrain tegmentum that has been described using multiple names and as containing multiple subnuclei by different neuroanatomists. In Olszewski and Baxter’s atlas, the most caudal aspect of the midbrain tegmentum, at the level of the isthmus, contains the cuneiform nucleus (CNF). The remainder of the caudal midbrain tegmentum, at the level of the decussation of the SCP, contains the mesencephalic reticular formation (MRF) and the mesencephalic reticular formation, ventral part (MRFv). Olszewski’s MRF nucleus continues rostrally and traverses the entire rostro-caudal extent of the midbrain, up to the mesencephalic-diencephalic junction. The MRFv’s rostral border ends at the mid-level of the red nuclei. In the rostral midbrain, at the level of the red nuclei, a small intracuneiform nucleus (ICUN) is shown within the MRF.

In Paxinos’ atlas, the most caudal aspect of the midbrain tegmentum, at the level of the isthmus, similarly contains CnF. However, another subnucleus named the isthmic reticular formation (isRt) is located in this region dorsal to the pedunculotegmental nucleus (PTg) and ventral to the CnF. In Paxinos’ atlas, the remainder of the caudal midbrain tegmentum, at the level of the decussation of the SCP, contains the CnF and isRt. At the transition from the decussation of the SCP to the red nuclei, the CnF and isRt nomenclature changes to the precuneiform area (PrCnF) and mesencephalic reticular formation (mRt), respectively. In the rostral midbrain, at the level of the red nuclei, the entire tegmentum is comprised of the mRt, which extends from the PAG on its dorso-medial border to the medial lemniscus on its ventro-lateral border. Within the rostral portion of the mRt, Paxinos defines a small subnucleus called the central mesencephalic nucleus (CeMe) in the same region that Olszewski defines the ICUN. Finally, at the mesencephalic-diencephalic junction, Paxinos’ mRt becomes the p1 reticular formation (p1Rt) (plates 8.62 to 8.64).

In summary, Olszewski and Paxinos describe a similar region of the midbrain tegmentum as containing a cluster of arousal nuclei, albeit with different nomenclature and different subnuclei. The region of the midbrain tegmentum that is described by Olszewski as containing CNF, MRF, MRFv, and ICUN is described by Paxinos as containing CnF, isRt, PrCnF, mRt, CeMe, and p1Rt. In our prior work (*9*), we used the term cuneiform/subcuneiform nucleus (CSC) to describe this region. We traced it in the rostral midbrain, in deference to Moruzzi and Magoun’s seminal study in which stimulation of the rostral midbrain tegmentum led to arousal and cortical activation in lightly anesthetized cats (*13*). We now update our nomenclature from CSC to mRt to ensure consistency with the Paxinos atlas’ nomenclature. Furthermore, we have extended our mRt region of interest (ROI) to include all of the aforementioned subnuclei described by Olszewski and Paxinos, such that our mRt ROI now extends throughout the entire rostro-caudal axis of the midbrain tegmentum. Specifically, our mRt ROI contains all of the following subnuclei described by Paxinos: CnF, isRt, PrCnF, mRt, CeMe, and p1Rt.

#### Periaqueductal Grey (PAG)

The anatomic borders and contours of our PAG ROI are consistent with those of the PAG described in Paxinos. We trace the PAG as a single ROI, which encompasses multiple subnuclei described in Paxinos. Specifically, our PAG ROI includes the following subnuclei described by Paxinos: lateral PAG (LPAG); dorsolateral PAG (DLPAG); dorsomedial PAG (DMPAG); pleioglia PAG (PlGl); and p1 PAG (p1PAG).

#### Pedunculotegmental Nucleus (PTg)

The PTg ROI contains both the pars compactus (called PTg by Paxinos) and the pars dissipatus (called the retroisthmic nucleus [RIs] by Paxinos). The pars dissipatus is called nucleus tegmenti pedunculopontinus, subnucleus dissipatus by Olszewski and the diffuse medial component of Ch5 by Mesulam (*196*). We have not changed our operational definition of neuroanatomic localization of the PTg, but we have updated our nomenclature from pedunculopontine nucleus (PPN) to PTg to ensure consistency with nomenclature of the Paxinos atlas. Our PTg ROI thus contains the PTg and RIs, as described by Paxinos. In addition, we made one edit to the boundaries of PTg in the Harvard AAN Atlas version 2.0, as compared to version 1.0: in version 2.0, we deleted one voxel in the PTg at MNI axial level z53 and reassigned this voxel to the VTA, based on tyrosine hydroxylase staining data and based on the anatomic boundaries in the Paxinos atlas.

#### Dorsal Raphe (DR)

The DR nucleus extends from the rostral pons to the caudal midbrain. We classify DR as a midbrain nucleus, because that is where most DR neurons are located. Paxinos shows DR continuing rostrally past the caudal border of the red nuclei. However, in our experience studying human brainstem specimens, serotonergic DR neurons do not extend past the rostral border of the trochlear nucleus, which is at the caudal border of the red nucleus. Our DR ROI includes the following subnuclei described by Paxinos: DR, caudal part (DRC); DR, dorsal part (DRD); DR, interfascicular part (DRI); DR, lateral part (DRL); DR, ventral part (DRV); posterodorsal raphe nucleus (PDR).

#### Ventral Tegmental Area (VTA)

The VTA ROI used in this study has two major differences compared to the prior VTA ROI used by our group and with version 1.0 of the Harvard Ascending Arousal Network Atlas: 1) its lateral extent and 2) its rostral extent. In our prior work, we did not include the lateral wing-like extensions of the PBP, as shown in Paxinos and in Pearson (*197*), because this subregion is difficult to differentiate from the nearby substantia nigra. However, with our new tyrosine hydroxylase immunostaining (Figure 1) and with reference to the Paxinos atlas, we were able to make this distinction between PBP and SN for the present studies.

The VTA node proposed here also has a more rostral extent than the VTA ROI that we used in prior studies (*9, 99*). Previously, we based the neuroanatomic borders of the VTA on hematoxylin-and-eosin-stained sections from human brainstem specimens, which revealed the highest density of catecholamine neurons, as demonstrated by the presence of neuromelanin, in the caudal midbrain. However, we recognize that VTA neurons have been identified in the rostral mesencephalon by other laboratories using a variety of staining techniques, such as tyrosine hydroxylase (*86, 198*). Thus, we performed new tyrosine hydroxylase stains on four midbrain sections from Specimen 1 (https://histopath.nmr.mgh.harvard.edu). These stains revealed a distribution of VTA neurons that was consistent with that described in the studies by Oades, Halliday, Pearson, and colleagues (**Figure 2**). Therefore, we use a VTA ROI in the present work that extends throughout the entire rostro-caudal axis of the midbrain. Finally, we do not include the rostral linear nucleus as part of the VTA, although some dopaminergic cells may reside in this region (*86*), because this nucleus is primarily serotonergic.

Finally, we edited the midline portion of the VTA node, based on the observation that VTA neurons in the midline of the midbrain were only seen at the level of the red nucleus, not at the level of the superior cerebellar peduncle. This immunostaining observation is consistent with the VTA anatomic borders in the Paxinos atlas along the rostro-caudal axis of the midbrain. Thus, we removed the midline VTA voxels at the level of the superior cerebellar peduncle in version 2.0 of the Harvard AAN atlas.

In summary, the VTA ROI used in this study contains the following nuclear subregions described by Paxinos: parainterfascicular nucleus (PIF); paranigral nucleus of ventral tegmental area (PN); ventral tegmental area (VTA); ventral tegmental area, rostral part (VTAR); and parabrachial pigmented nucleus of the VTA (PBP). This VTA ROI contains the A10 nuclei, as described by Oades and Halliday (*86*). The neuroanatomic borders of the VTA ROI contain the following subregions described by Olszewski: the paranigral nucleus (PNg), interfascicular nucleus (INF), parabrachial pigmented nucleus (PBP), and ventral tegmental area of Tsai (VTA).

### Neuroanatomic localization of pontine dAAN nodes

#### Pontine Reticular Nucleus, Oral Part (i.e., Pontis Oralis; PnO)

We have not changed the neuroanatomic localization of this nucleus since our prior studies (*9*), but we have updated our nomenclature from PO to PnO to reflect the Paxinos atlas’s nomenclature. Our PnO nucleus includes the following subnuclei described by Olszewski: nucleus reticularis pontis oralis (NRPO) and nucleus reticularis tegmenti pontis (NRTP). Paxinos does not subdivide PnO into these two subnuclei.

#### Parabrachial Complex (PBC)

The PBC is divided in the Paxinos atlas into medial and lateral components: medial parabrachial nucleus (MPB) and lateral parabrachial nucleus (LPB). MPB and LPB are further subdivided into subregions: MPB, external part (MPBE); LPB, central part (LPBC); LPB, dorsal part (LPBD); LPB, external part (LPBE); LPB, superior part (LPBS). Given the difficulty of delineating these subregions on MRI scans of the human brain, even at the ultra-high spatial resolution (∼600 µm) at which Specimen 1 was imaged, we grouped all of these medial and lateral subregions into a single PBC node. This is the same approach that we used to delineate the PBC in our previous analyses of Specimen 1.

Of note, the caudal boundary of PBC in Paxinos (MPB) is located at plate 8.35, but we operationally define the caudal boundary to be located at the level of plate 8.37, which is the caudal border of the pontine arousal nuclei. Similarly, the rostral boundary of PBC in Paxinos’ atlas is located at plate 8.49 but we operationally define the PBC as extending to plate 8.50, which is the transition point between PBC and PTg at the ponto-mesencephalic junction.

#### Median Raphe Nucleus (MnR)

In accordance with the terminology used by Paxinos, we define the MnR as comprising both the MnR proper and the slightly more lateral subnucleus, the paramedian raphe nucleus (PMnR). The PMnR begins in the midpons on plate 8.38, ventral to the dorsal raphe and medial to the nucleus pontis oralis. It is joined medially by the MnR on plate 8.44 and disappears prior to the isthmus, on plate 8.48. The MnR continues rostrally into the caudal midbrain, ending at the level of the decussation of the superior cerebellar peduncle. As with the PBC (above), we operationally define the caudal and rostral boundaries of our MnR ROI as plates 8.37 and 8.50, respectively, since these are the anatomic levels of the mid-pons and the ponto-mesencephalic junction. Our MnR ROI encompasses the MnR and paramedian raphe nucleus.

#### Locus Coeruleus (LC)

This small, tubular nucleus begins in the dorsolateral midpons, on Paxinos plate 8.37, lateral to the fourth ventricle and medial to the superior cerebellar peduncle. It extends rostrally to the isthmus on plate 8.49. As with PBC and MnR above, we operationally consider the LC as extending to 8.50, which is the ponto-mesencephalic junction. The neuroanatomic location, borders and nomenclature for LC are similar in the Olszewski and Paxinos atlases.

#### Laterodorsal Tegmental Nucleus (LDTg)

Using Paxinos’ terminology, we do not include the ventral nucleus, which is labeled the laterodorsal tegmental nucleus, ventral part (LDTgV), as part of our annotation of the LDTg. The LDTg is immediately adjacent to the medial edge of the LC. It lies dorsal to the medial longitudinal fasiculus (mLF), while the LDTgV is the subarea ventral to the mLF. It begins several levels rostral to the mid-pons (plate 8.41) and terminates at the level of the trochlear nucleus (8.51). As with PBC, MnR, and LC above, we operationally consider the LDTg as extending to 8.50, which is the ponto-mesencephalic junction.

### Neuroanatomic localization of thalamic dAAN nodes

#### Intralaminar Nuclei (ILN)

Using the terminology of the Ding Brain Atlas (*49*), this collection of thalamic subnuclei includes the central lateral (CL), central medial (CeM), paracentral (PC), central dorsal (CD), centromedian (CM), parafascicular (Pf), subparafasicular (SPf) and the fasiculosus (Fa) nuclei. These subnuclei form an arc with a ventro-posterior bulge and are bordered medially by the dorsomedial nucleus and laterally by ventral anterior nucleus, ventral posterior nucleus and pulvinar.

#### Reticular Nucleus (Ret)

The reticular thalamic nucleus includes the magnocellular (Rmc) and parvocellular (Rpc) divisions (*49*) and forms the anterior and lateral border of the thalamus. Posteriorly, at the level of the lateral geniculate nucleus, it is laterally separated from the pulvinar by a thin layer of white matter.

#### Paraventricular Thalamic Nucleus (PaV)

The paraventricular thalamic nucleus (PaV) lines the medial border of the thalamus. On its medial side is the third ventricle. On its lateral side are several different thalamic nuclei along its antero-posterior extent: anteriorly, the lateral border of PaV is formed by ILN and the anterior nuclear complex; posteriorly, the lateral border of PaV is formed by the mediodorsal nucleus of the thalamus. The superior border of PaV is formed by the stria medullaris of the thalamus, and the inferior border of PaV is formed by the midline nuclear complex.

### Neuroanatomic localization of hypothalamic dAAN nodes

#### Lateral Hypothalamic Area (LHA)

This region is subdivided into anterior (LHAa), posterior (LHAp) and tuberal regions (LHAtub) and forms the lateral border of the hypothalamus with the substantia innominata of the basal forebrain (*49*). The anterior-most region is not depicted in the Allen atlas. The tuberal region begins posterior to the ventrolateral pre-optic area (VLPO) and is subdivided into a magnocellular nucleus (LHmc), accessory secretory cells (LHsc) and a pallidohypothalamic area (PalHy). The posterior region extends to the posterior border of the hypothalamus, dorsal to the mammillary bodies and includes the perifornical nucleus (PeF).

#### Tuberomammillary Nucleus (TM)

This nucleus has an anterior component within the LHAtub, just medial to the optic tract. Posteriorly, it sits laterally to the mammillary bodies and inferior to the LHAp.

#### Supramammillary Nucleus (SUM)

This nucleus is located superior to the mammillary body and mammillothalamic tract. It is bordered medially by the posterior hypothalamic nucleus and laterally by the LHA and the TMN.

### Neuroanatomic localization of basal forebrain dAAN nodes

#### Basal Nucleus of Meynert/Substantia Innominata (BNM/SI)

The BNM and SI are identified as a single region in our studies, as they cannot be anatomically distinguished using MRI. They occupy the area lateral to the hypothalamus and ventral to the pallidum, extending anteriorly to the level of the anterior commissure and posteriorly to the posterior boundary of the amygdala (*49*).

#### Nucleus of the Diagonal Band (NDB)

The NDB includes vertical and horizontal subdivisions. It exists in its vertical orientation anterior to the anterior commissure, in the midline, medial to the septal nucleus and nucleus accumbens. At the level of the anterior commissure, it condenses ventrally into its horizontal subdivision, forming the border between the hypothalamus and the SI (*49*).

### Methodologic Considerations for Deterministic and Probabilistic Tractography

To ensure consistency across methods, we standardized the deterministic and probabilistic processing parameters wherever possible, as detailed in the main text. Some processing parameters, however, were not standardized due to differences in the deterministic and probabilistic techniques. For example, in all deterministic experiments, fiber tracts were terminated when the angle between water diffusion vectors in adjacent voxels exceeded a threshold of 60°. This angle threshold was previously identified by our group as providing a balance between tract sensitivity (i.e. identification of the brainstem’s widely branching pathways) and tract specificity (i.e. reduction of false positive tracts) (*9, 54, 99*). In contrast, we and other groups typically use a more liberal angle threshold of 80° (curvature threshold = 0.2), for probabilistic experiments, consistent with the default setting in FSL’s Diffusion Toolbox. Notably, a recent *ex vivo* probabilistic tractography study that compared the accuracy of fiber tract data in the macaque brain to “gold standard” tract-tracer data found that probabilistic tract accuracy was minimally affected when angle thresholds were in the range of 70-90° (*186*). Similarly, the accuracy of the probabilistic tractography data was not affected by varying the FA thresholds between 0.0-0.3.

We did not include a distance correction (e.g., the “-pd” function in FSL’s probtrackx) in our *CP* calculations. Although the probability of a connection between two anatomic regions, as detected by tractography, may be associated with the distance between the regions (*199*), it is unknown whether this association is linear or exponential. Moreover, the type of distance correction that is appropriate for a given bundle of tracts likely depends upon multiple variables, including the number of voxels in the seed/target nodes, the number of other bundles that intersect the bundle of interest along its course, and the geometry of the bundle (e.g. its curvature and branching). From a statistical perspective, it is unclear if a single distance correction method can account for the effect of distance on connectivity measures for different bundles. Furthermore, empiric data from a recent *ex vivo* MRI study of the macaque brain, which tested linear and exponential distance correction methods, found that the accuracy of probabilistic tractography data against gold-standard tract-tracing data was not improved by either distance correction method (*186*). Thus, we did not correct for inter-nodal distance in our *CP* measurements.

## Supplementary Videos

**Video S1. The Harvard Ascending Arousal Network (AAN) Atlas – Version 2.0.** AAN brainstem nodes are rendered in three dimensions and superimposed upon the MNI152 atlas in 1 mm isotropic space. For anatomic reference, the nodes are shown on an axial image at the level of the mid-pons, a sagittal image at the midline, and a coronal image at the splenium of the corpus callosum. Blue = locus coeruleus; yellow = parabrachial complex; green = median raphe; orange = laterodorsal tegmental nucleus; light blue = pontis oralis; turquoise = dorsal raphe; red = mesencephalic reticular formation; purple = pedunculotegmental nucleus; light purple = periaqueductal grey; pink = ventral tegmental area.

**Video S2. Visualizing default ascending arousal network connectivity within the ventral tegmental area hub node.** All candidate brainstem dAAN nodes and their tract end-points are shown from a dorsal perspective, as in Fig. 6, superimposed upon a coronal non-diffusion-weighted (*b=0*) image at the level of the mid-thalamus and an axial *b=0* image at the intercollicular level of the midbrain. Tract end-points appear as disks and represent the start-points and termination-points of each tract (i.e., there are two end-points per tract). All tract end-points are color-coded by the node from which they originate, and the nodes are rendered semi-transparent so that end-points can be seen within them.

As the video begins, tract end-points are seen within the semi-transparent brainstem candidate nodes: ventral tegmental area (VTA), periaqueductal grey (PAG), and mesencephalic reticular formation (mRt), pedunculotegmental nucleus (PTg), and parabrachial complex (PBC). As the video proceeds, the viewer’s eye moves ventrally through the PAG. Once the viewer’s eye emerges from the ventral surface of the PAG, the dorsomedial border of the VTA is seen, where tract end-points from multiple candidate brainstem dAAN nodes overlap within the VTA and along its dorsal border: pedunculotegmental, parabrachial complex, and dorsal raphe (PTg-PBC-DR); pontis oralis, DR, and PTg (PnO-DR-PTg); and PBC-mRt-PTg. Tract end-point overlap does not prove synaptic connectivity but indicates extensive connectivity via association pathways between ipsilateral and midline dAAN nodes with the VTA.

## Supplementary Figures

**Fig. S1.**
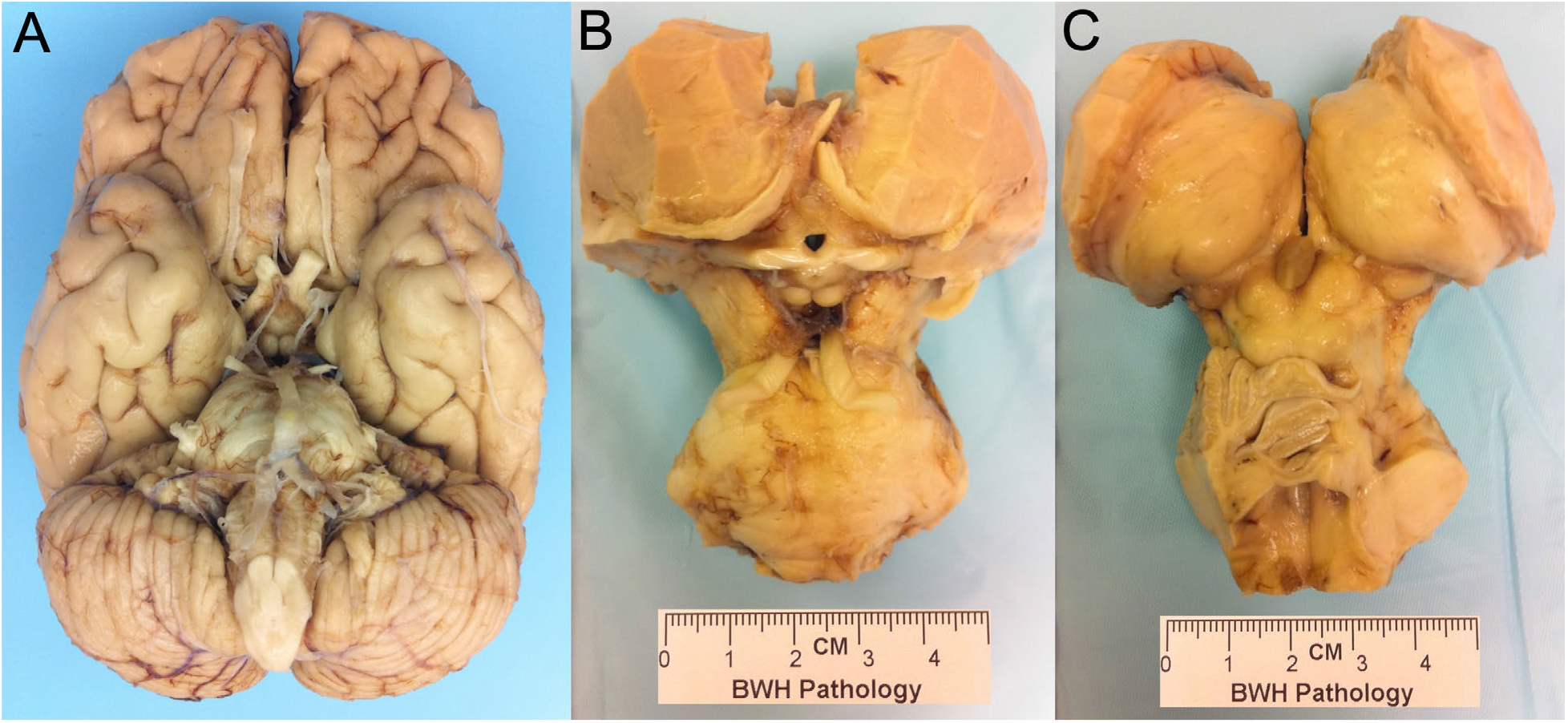
Gross pathology photos of Specimen 1. (**A**) Inferior view of brain Specimen 1. The whole brain was dissected to a brainstem/diencephalon/forebrain specimen consisting of the pons, midbrain, hypothalamus, thalamus, basal forebrain, and portions of the basal ganglia, as shown from an (**B**) anterior and (**C**) posterior perspective. CM = centimeters. Figure adapted from Edlow et al. JNEN 2012 (*9*).

**Fig. S2.**
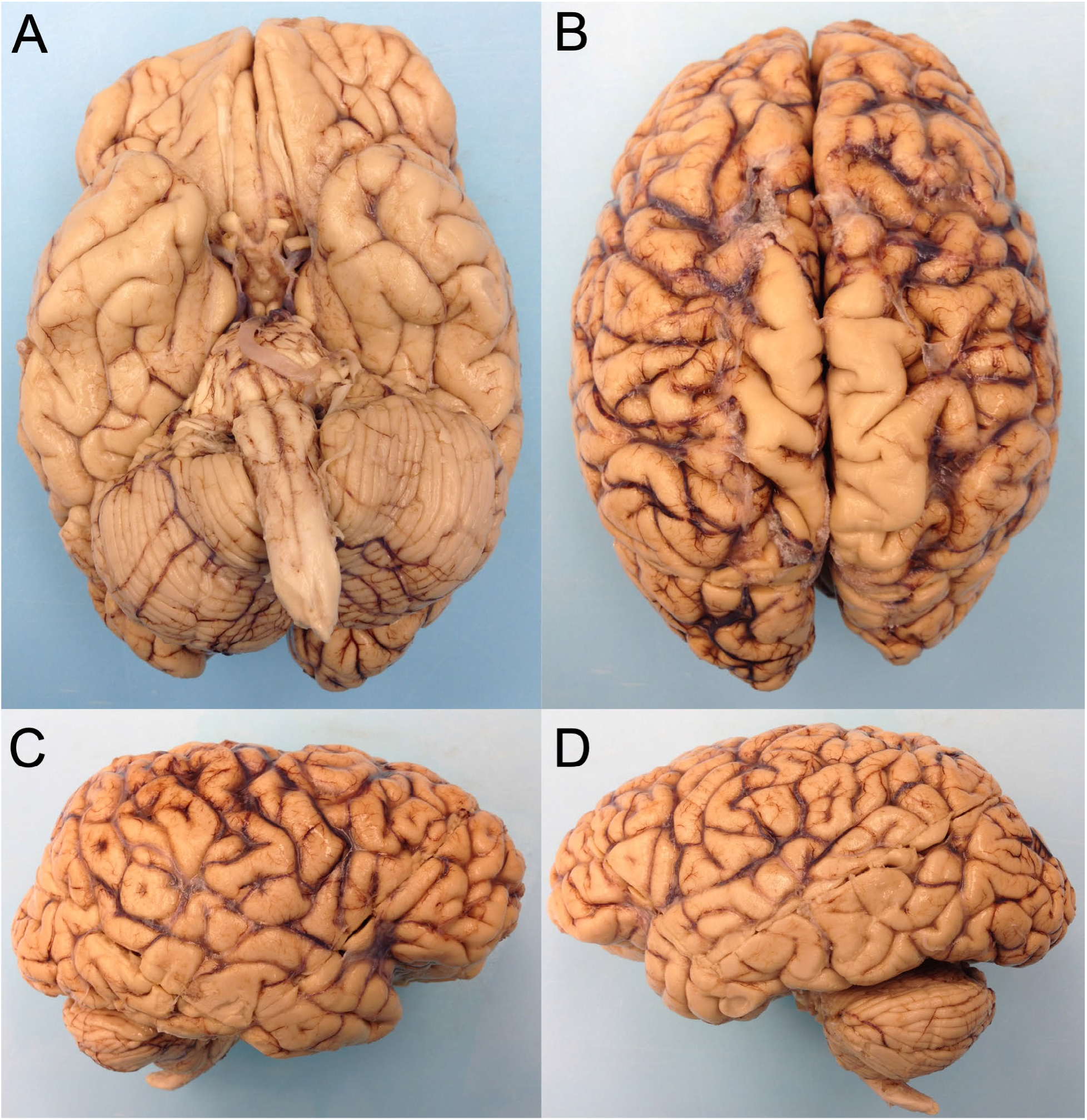
Gross pathology photos of Specimen 2. Brain Specimen 2 is shown from an (**A**) inferior, (**B**) superior, (**C**) right lateral, and (**D**) left lateral perspective.

**Fig. S3.**
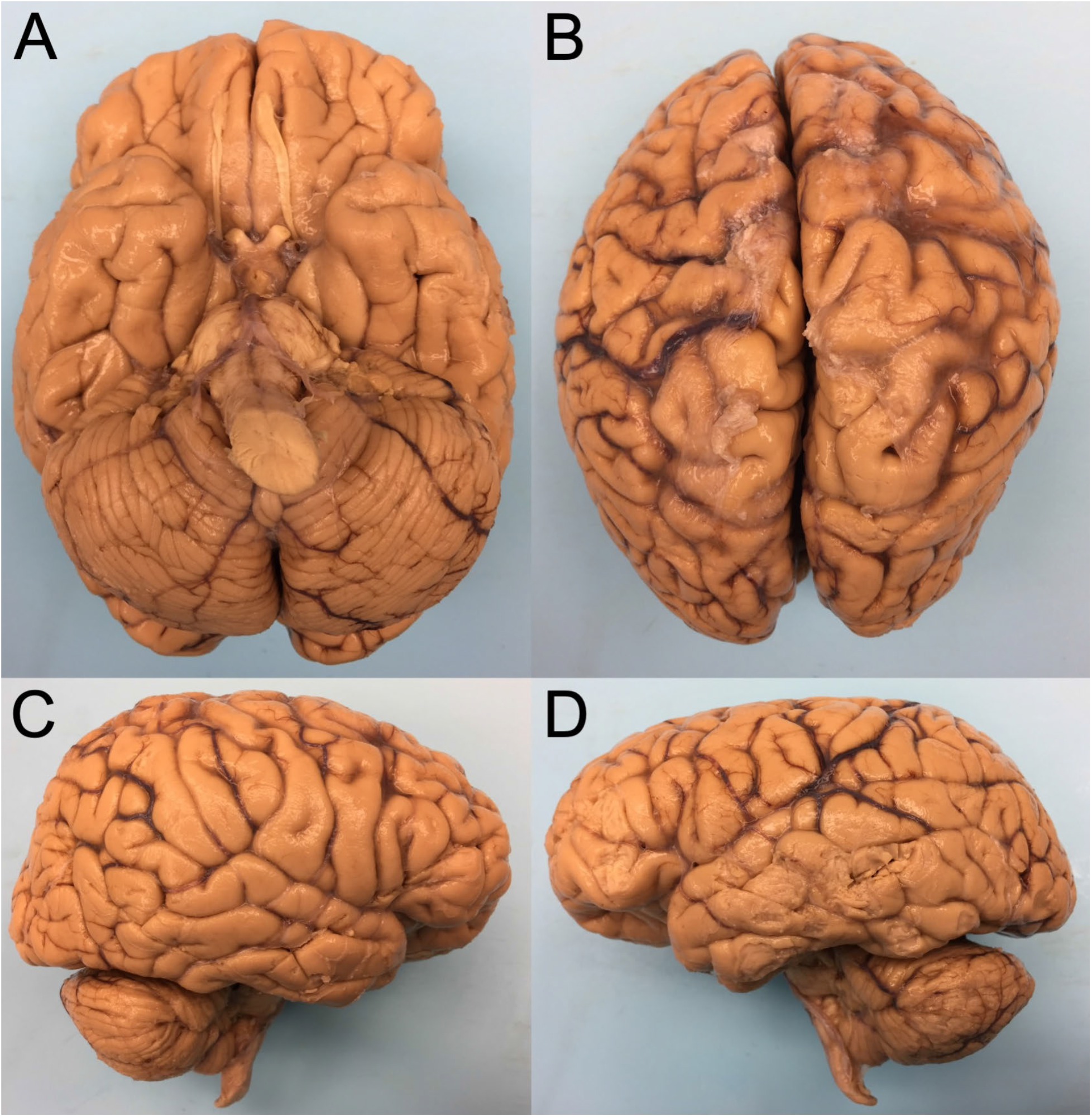
Gross pathology photos of Specimen 3. Brain Specimen 3 is shown from an (**A**) inferior, (**B**) superior, (**C**) right lateral, and (**D**) left lateral perspective.

**Fig S4.**
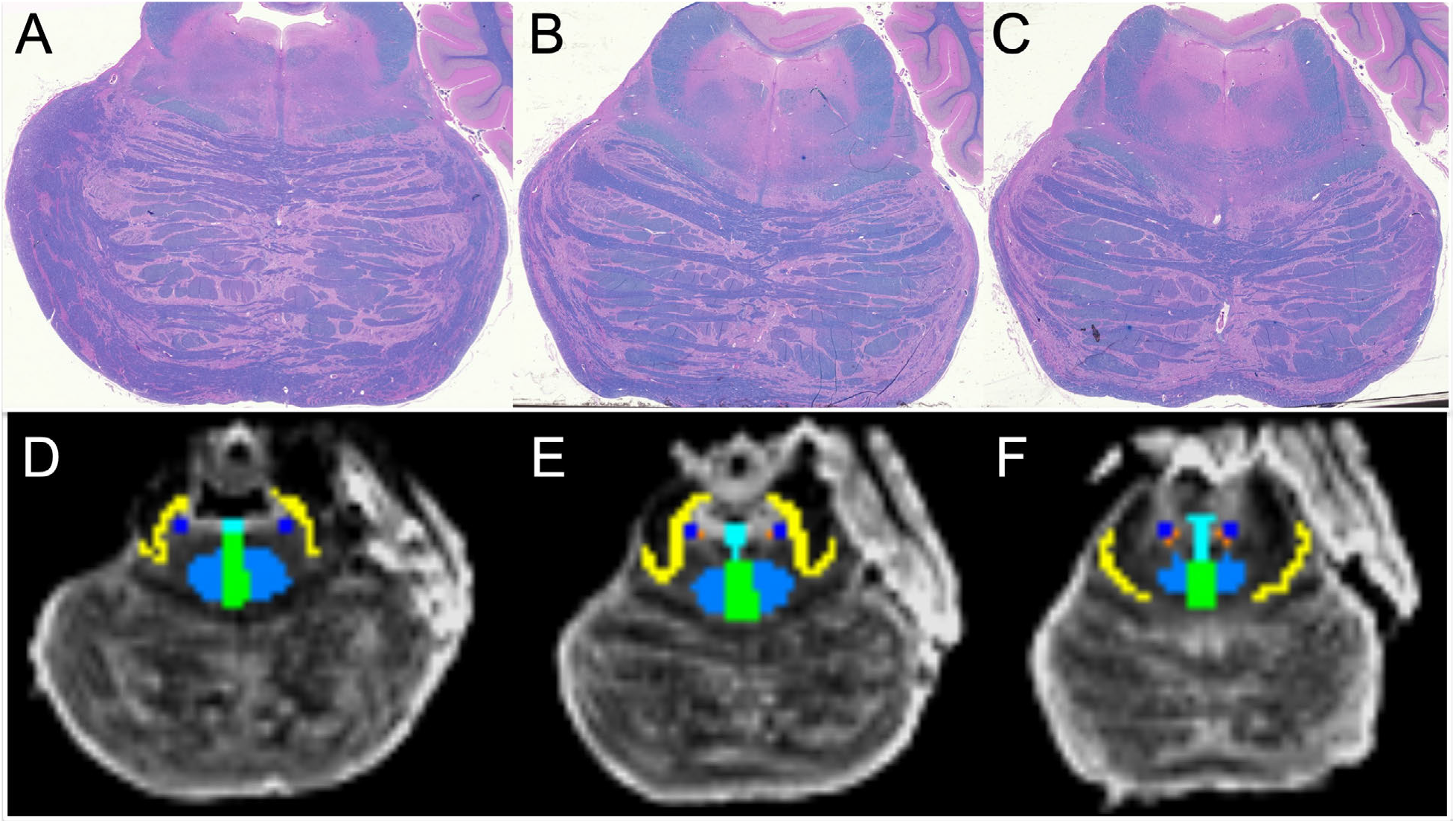
Neuroanatomic Localization of Pontine Arousal Nuclei. Axial histologic sections through the mid pons (**A**), upper pons (**B**) and rostral pons (**C**) for Specimen 1. Each section is stained with hematoxylin and eosin/Luxol fast blue. Corresponding axial non-diffusion-weighted (*b=0*) images (**D, E, F**). Regions of interest for tractography are traced on the *b=0* images with neuroanatomic borders determined by the histological stains: light blue = pontis oralis; turquoise = dorsal raphe; dark blue = locus coeruleus; yellow = parabrachial complex; green = median raphe; orange = laterodorsal tegmental nucleus. Figure adapted from Edlow et al. JNEN 2012 (*9*).

**Fig. S5.**
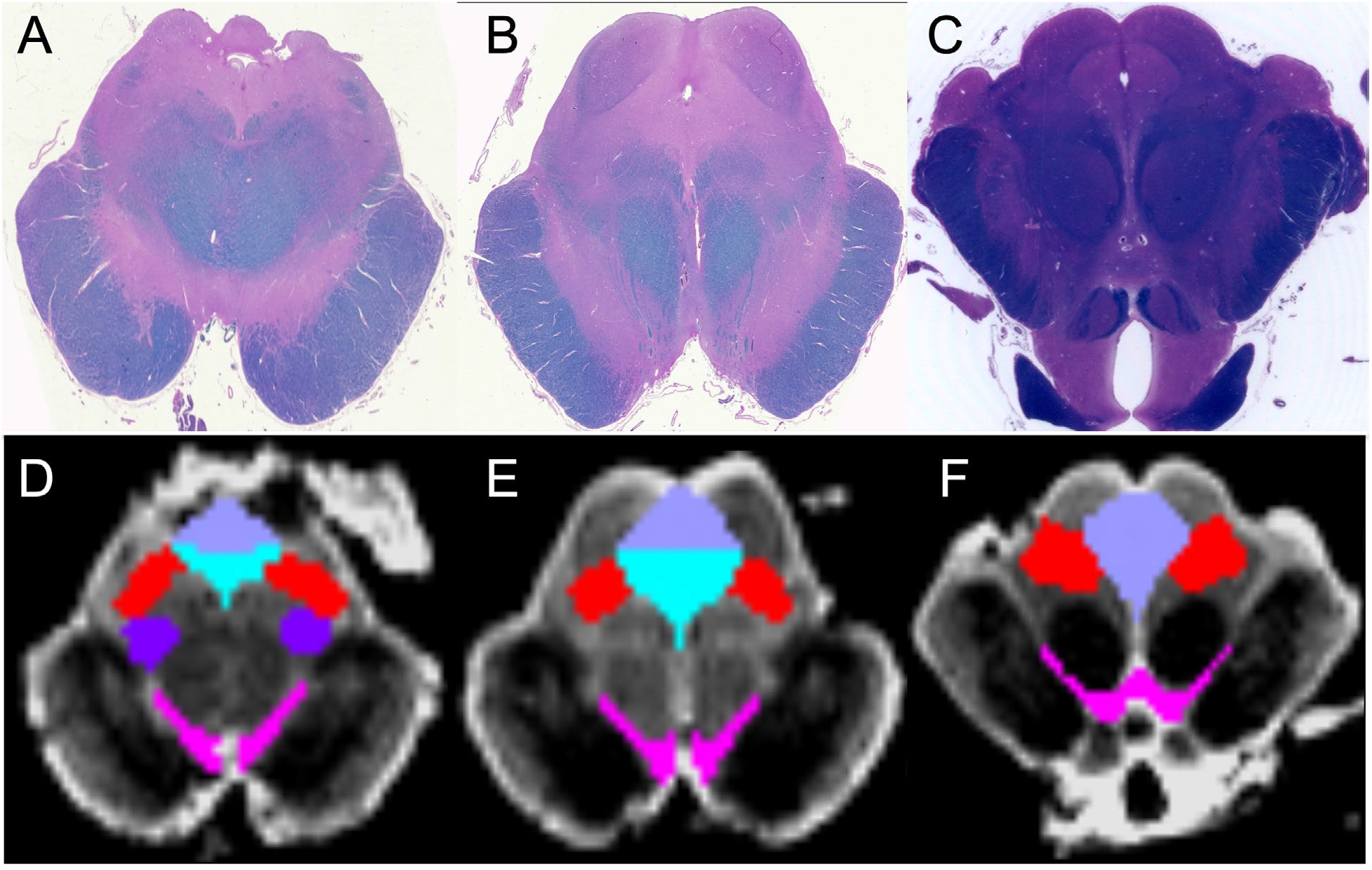
Neuroanatomic Localization of Midbrain Arousal Nuclei. Axial histologic sections through the midbrain at the level of the decussation of the superior cerebellar peduncles (**A**), at the level of the inferior colliculi (**B**) and at the level of the superior colliculi and red nuclei (**C**) in Specimen 1. All sections are stained with hematoxylin and eosin/Luxol fast blue. Corresponding axial non-diffusion-weighted (*b=0*) images (**D, E, F**). Regions of interest for tractography are traced on the *b=0* images with neuroanatomic borders determined by the histological stains: red = mesencephalic reticular formation; pink = ventral tegmental area; dark purple = pedunculotegmental nucleus; turquoise = dorsal raphe; light purple = periaqueductal grey. Figure adapted from Edlow et al. JNEN 2012 (*9*).

## Supplementary Tables

**Table S1.**
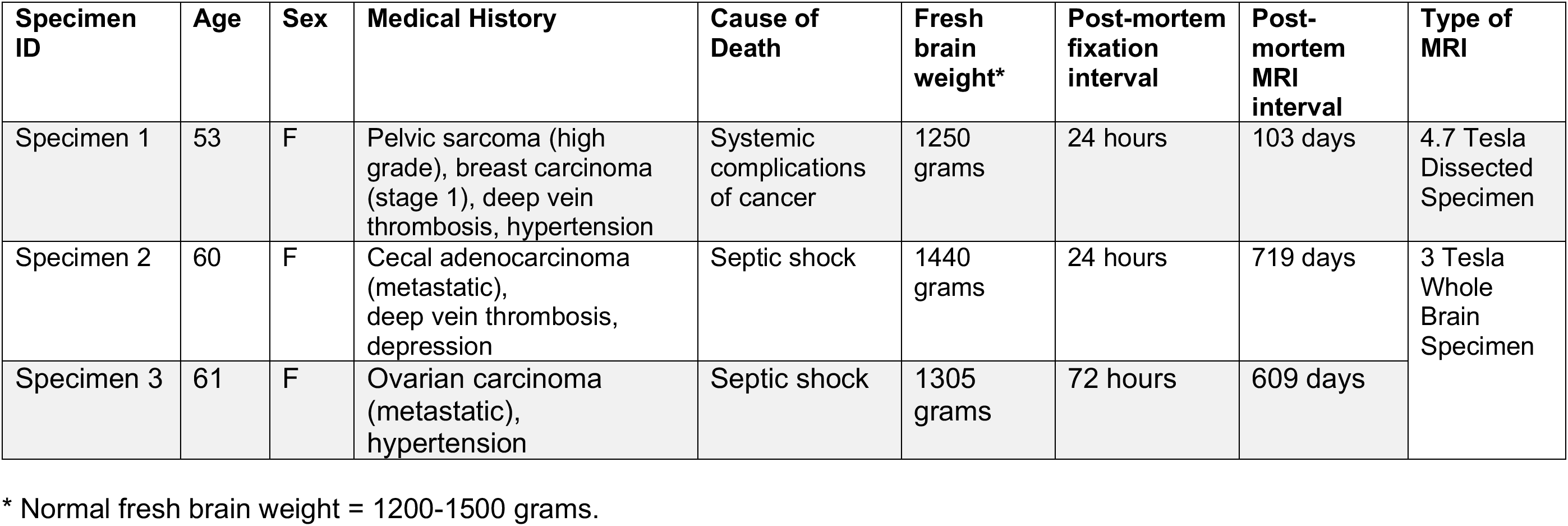
Demographic and clinical characteristics for individuals who donated brain specimens.

**Table S2.**
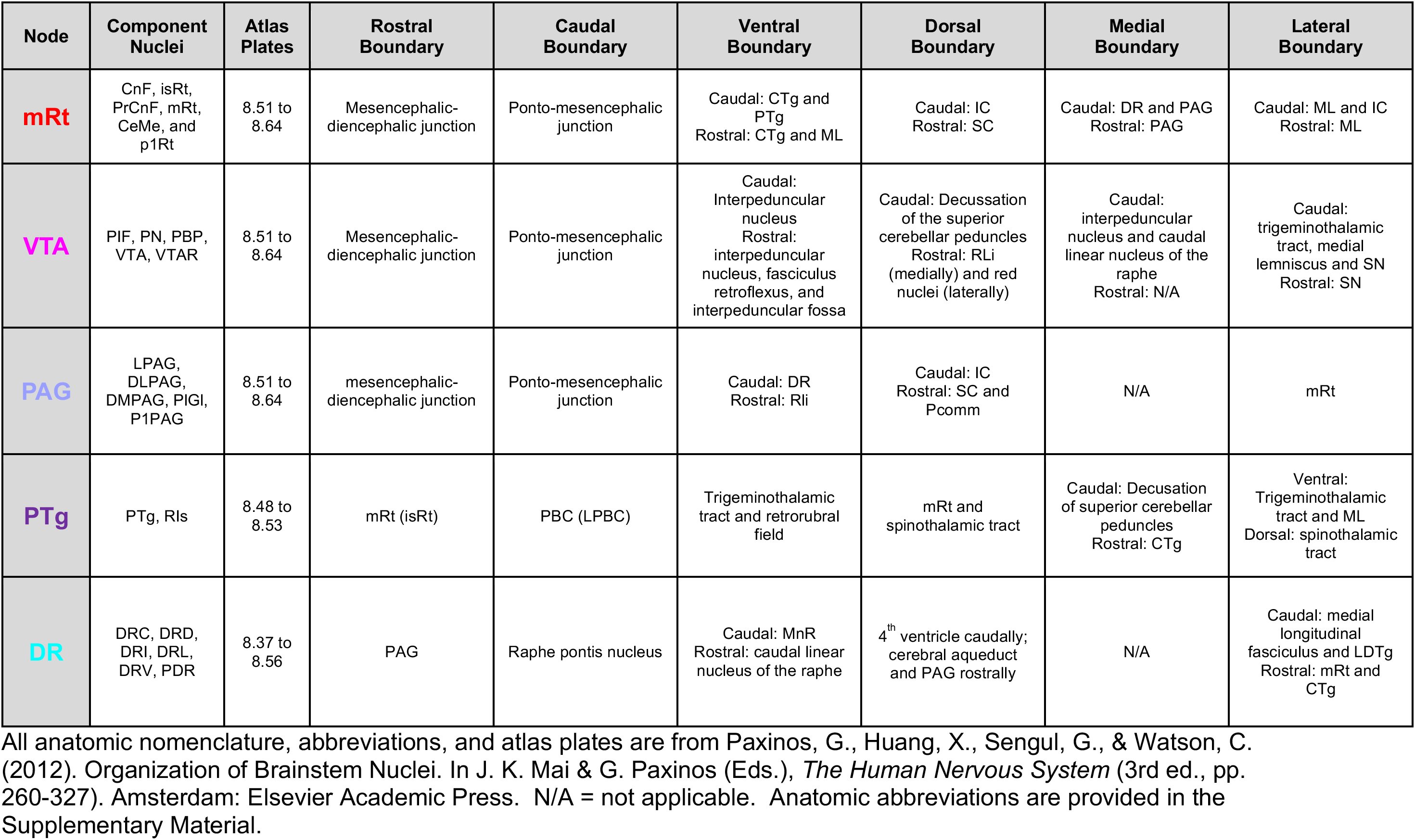
Neuroanatomic Localization of Midbrain Arousal Nuclei.

| Node | Component Nuclei | Atlas Plates | Rostral Boundary | Caudal Boundary | Ventral Boundary | Dorsal Boundary | Medial Boundary | Lateral Boundary |
| --- | --- | --- | --- | --- | --- | --- | --- | --- |
| **mRt** | CnF, isRt, PrCnF, mRt, CeMe, and p1Rt | 8.51 to 8.64 | Mesencephalic-diencephalic junction | Ponto-mesencephalic junction | Caudal: CTg and PTg<br>Rostral: CTg and ML | Caudal: IC<br>Rostral: SC | Caudal: DR and PAG<br>Rostral: PAG | Caudal: ML and IC<br>Rostral: ML |
| **VTA** | PIF, PN, PBP, VTA, VTAR | 8.51 to 8.64 | Mesencephalic-diencephalic junction | Ponto-mesencephalic junction | Caudal: Interpeduncular nucleus<br>Rostral: interpeduncular nucleus, fasciculus retroflexus, and interpeduncular fossa | Caudal: Decussation of the superior cerebellar peduncles<br>Rostral: RLi (medially) and red nuclei (laterally) | Caudal: interpeduncular nucleus and caudal linear nucleus of the raphe<br>Rostral: N/A | Caudal: trigeminothalamic tract, medial lemniscus and SN<br>Rostral: SN |
| **PAG** | LPAG, DLPAG, DMPAG, PIGI, P1PAG | 8.51 to 8.64 | mesencephalic-diencephalic junction | Ponto-mesencephalic junction | Caudal: DR<br>Rostral: Rli | Caudal: IC<br>Rostral: SC and Pcomm | N/A | mRt |
| **PTg** | PTg, Rls | 8.48 to 8.53 | mRt (isRt) | PBC (LPBC) | Trigeminothalamic tract and retrorubral field | mRt and spinothalamic tract | Caudal: Decussation of superior cerebellar peduncles<br>Rostral: CTg | Ventral: Trigeminothalamic tract and ML<br>Dorsal: spinothalamic tract |
| **DR** | DRC, DRD, DRI, DRL, DRV, PDR | 8.37 to 8.56 | PAG | Raphe pontis nucleus | Caudal: MnR<br>Rostral: caudal linear nucleus of the raphe | 4<sup>th</sup> ventricle caudally; cerebral aqueduct and PAG rostrally | N/A | Caudal: medial longitudinal fasciculus and LDTg<br>Rostral: mRt and CTg |

All anatomic nomenclature, abbreviations, and atlas plates are from Paxinos, G., Huang, X., Sengul, G., & Watson, C. (2012). Organization of Brainstem Nuclei. In J. K. Mai & G. Paxinos (Eds.), *The Human Nervous System* (3rd ed., pp. 260-327). Amsterdam: Elsevier Academic Press. N/A = not applicable. Anatomic abbreviations are provided in the Supplementary Material.

**Table S3.**
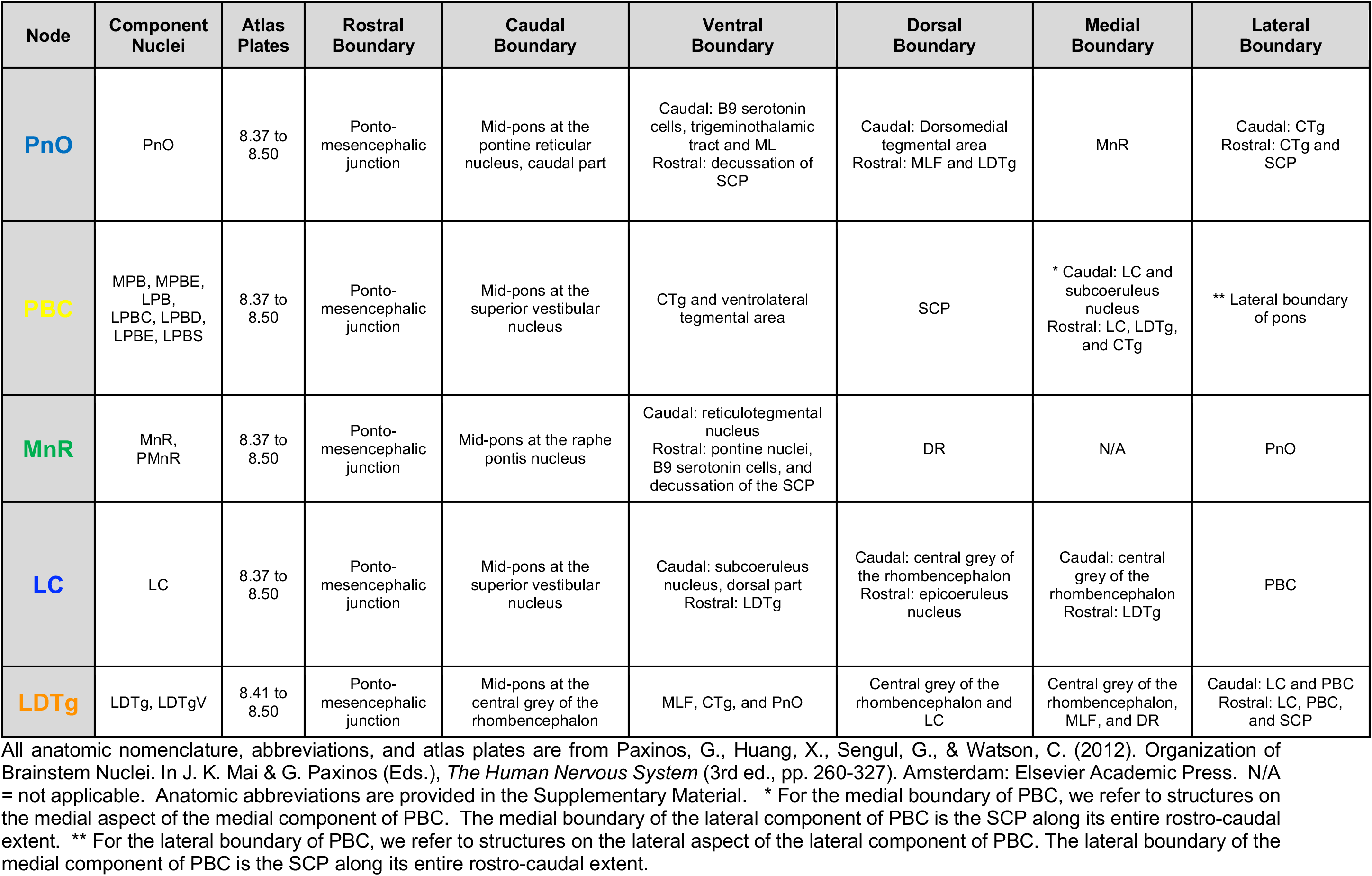
Neuroanatomic Localization of Pontine Arousal Nuclei.

| Node | Component Nuclei | Atlas Plates | Rostral Boundary | Caudal Boundary | Ventral Boundary | Dorsal Boundary | Medial Boundary | Lateral Boundary |
| --- | --- | --- | --- | --- | --- | --- | --- | --- |
| **PnO** | PnO | 8.37 to 8.50 | Ponto-mesencephalic junction | Mid-pons at the pontine reticular nucleus, caudal part | Caudal: B9 serotonin cells, trigeminothalamic tract and ML<br>Rostral: decussation of SCP | Caudal: Dorsomedial tegmental area<br>Rostral: MLF and LDTg | MnR | Caudal: CTg<br>Rostral: CTg and SCP |
| **PBC** | MPB, MPBE, LPB, LPBC, LPBD, LPBE, LPBS | 8.37 to 8.50 | Ponto-mesencephalic junction | Mid-pons at the superior vestibular nucleus | CTg and ventrolateral tegmental area | SCP | * Caudal: LC and subcoeruleus nucleus<br>Rostral: LC, LDTg, and CTg | ** Lateral boundary of pons |
| **MnR** | MnR, PMnR | 8.37 to 8.50 | Ponto-mesencephalic junction | Mid-pons at the raphe pontis nucleus | Caudal: reticulotegmental nucleus<br>Rostral: pontine nuclei, B9 serotonin cells, and decussation of the SCP | DR | N/A | PnO |
| **LC** | LC | 8.37 to 8.50 | Ponto-mesencephalic junction | Mid-pons at the superior vestibular nucleus | Caudal: subcoeruleus nucleus, dorsal part<br>Rostral: LDTg | Caudal: central grey of the rhombencephalon<br>Rostral: epicoeruleus nucleus | Caudal: central grey of the rhombencephalon<br>Rostral: LDTg | PBC |
| **LDTg** | LDTg, LDTgV | 8.41 to 8.50 | Ponto-mesencephalic junction | Mid-pons at the central grey of the rhombencephalon | MLF, CTg, and PnO | Central grey of the rhombencephalon and LC | Central grey of the rhombencephalon, MLF, and DR | Caudal: LC and PBC<br>Rostral: LC, PBC, and SCP |

All anatomic nomenclature, abbreviations, and atlas plates are from Paxinos, G., Huang, X., Sengul, G., & Watson, C. (2012). Organization of Brainstem Nuclei. In J. K. Mai & G. Paxinos (Eds.), *The Human Nervous System* (3rd ed., pp. 260-327). Amsterdam: Elsevier Academic Press. N/A = not applicable. Anatomic abbreviations are provided in the Supplementary Material. \* For the medial boundary of PBC, we refer to structures on the medial aspect of the medial component of PBC. The medial boundary of the lateral component of PBC is the SCP along its entire rostro-caudal extent. \*\* For the lateral boundary of PBC, we refer to structures on the lateral aspect of the lateral component of PBC. The lateral boundary of the medial component of PBC is the SCP along its entire rostro-caudal extent.

**Table S4.** Neuroanatomic Localization of Thalamic Arousal Nuclei.

| Node | Component Nuclei | Atlas Plates | Inferior Boundary | Superior Boundary | Anterior Boundary | Posterior Boundary | Medial Boundary | Lateral Boundary |
| --- | --- | --- | --- | --- | --- | --- | --- | --- |
| **IL** | CL, CeM, PC, CD, CM, Pf, SPf, Fa | 27 to 57 | Centromedian nucleus of the thalamus | Anterior: anteroventral nucleus of the thalamus<br>Posterior: dorsal superficial nucleus (laterodorsal nucleus) of the thalamus | Anterior border of thalamus<br>Fasciculosus nucleus | Pulvinar | Medial dorsal thalamic nucleus | Pulvinar ventral lateral thalamic nucleus posteriorly and ventral anterior thalamic nucleus anteriorly |
| **Ret** | Rmc, Rpc | 25 to 65 | Zona incerta and subthalamic nucleus | Stria terminalis and white matter of the forebrain | Bed nucleus of stria terminalis | Crus of the fornix | Anterior: Ventral anterior nucleus of thalamus<br>Posterior: Ventral lateral nucleus, lateral posterior nucleus, and pulvinar | White matter of the forebrain (internal capsule) |
| **PaV** | PaVr, PaVc | 27 to 47 | Anterior: rhomboid nucleus of thalamus<br>Posterior: periventricular area of thalamus, intermediodorsal nucleus of thalamus, and intralaminar nuclei | Stria medullaris and velum interpositum | Supraoptic region of hypothalamus and fornix | Periventricular area of thalamus | Third ventricle | Anterior: fasciculosus nucleus of thalamus and anteromedial nucleus of thalamus<br>Posterior: mediodorsal nucleus and intralaminar nuclei |

All anatomic nomenclature, abbreviations, and atlas plates are from Ding, S.L. et al. (2016). Comprehensive cellular-resolution atlas of the adult human brain. *J Comp Neurol* **524** (16): 3127-3481. Anatomic abbreviations are provided in the Supplementary Material.

**Table S5.** Neuroanatomic Localization of Hypothalamic Arousal Nuclei.

| Node | Component Nuclei | Atlas Plates | Inferior Boundary | Superior Boundary | Anterior Boundary | Posterior Boundary | Medial Boundary | Lateral Boundary |
| --- | --- | --- | --- | --- | --- | --- | --- | --- |
| **TMN** | TMN | 34* to 40 | Lateral tuberal nuclei and medial mammillary nucleus | Lateral hypothalamic area and substantia nigra, reticular part | Lateral hypothalamic area | Lateral hypothalamic area and retromammillary area | Anterior: lateral hypothalamic area and fornix<br>Posterior: medial mammillary nucleus | Anterior: optic tract<br>Posterior: substantia nigra, reticular part |
| **LHA** | LHAa**, LHAp, LHAtub | 27 to 42 | Anterior: optic tracts, supraoptic nucleus, retrochiasmatic nucleus, and interpeduncular cistern<br>Posterior: TM and mammillary bodies | Anterior: dorsomedial hypothalamic nucleus, tuberal region of hypothalamus, and fornix<br>Posterior: ventral anterior nucleus of thalamus, mammillo-tegmental tract and zona incerta | Preoptic region of hypothalamus | Midbrain | Anterior: tuberal region of hypothalamus<br>Posterior: mammillary region of hypothalamus, mammillothalamic tract | Anterior: BNM/SI, forebrain white matter<br>Posterior: Subthalamic nucleus and substantia nigra reticular part |
| **SUM** | SUM | 35 to 40 | Mammillary body and mammillothalamic tract | Posterior hypothalamic nucleus | Posterior hypothalamic nucleus and mammillary peduncle | Posterior hypothalamic nucleus | Posterior hypothalamic nucleus | Anterior: LHA<br>Posterior: TMN and mammillary body |

All anatomic nomenclature, abbreviations, and atlas plates are from Ding, S.L. et al. (2016). Comprehensive cellular-resolution atlas of the adult human brain. *J Comp Neurol* **524** (16): 3127-3481. N/A = not applicable. Anatomic abbreviations are provided in the Supplementary Material.
\* From plates 29 to 33, TM is fully enveloped within LHA and cannot be reliably distinguished from LHA. Therefore, we operationally define the anterior boundary of the TM node at plate 34. \*\* LHAa is not depicted in the Ding atlas.

**Table S6.** Neuroanatomic Localization of Basal Forebrain Arousal Nuclei.

| Node | Component Nuclei | Atlas Plates | Inferior Boundary | Superior Boundary | Anterior Boundary | Posterior Boundary | Medial Boundary | Lateral Boundary |
| --- | --- | --- | --- | --- | --- | --- | --- | --- |
| **DBB** | NDBv, NDBh, dib | 19 to 23 | Anterior: subgenual division of anterior cingulate cortex<br>Posterior: basal cisterns | Anterior: medial septal nucleus<br>Posterior: preoptic region of hypothalamus | subgenual division of anterior cingulate cortex | Preoptic region of hypothalamus | Anterior: basal cisterns<br>Posterior: preoptic region of hypothalamus | Anterior: septal nuclei<br>Posterior: NBM/SI |
| **BNM/SI** | BNMI, BNMm, SI-pc, SI-nc | 19 to 45 | Anterior: forebrain white matter, subgenual division of anterior cingulate cortex<br>Middle: basal cisterns<br>Posterior: amygdaloid complex | Anterior: ventral pallidus, olfactory tubercle<br>Middle: sublenticular extended amygdala, anterior commissure<br>Posterior: globus pallidus | Olfactory tubercle, forebrain white matter | Optic tract, amygdaloid complex, forebrain white matter, | Anterior: diagonal band<br>Middle: LHA<br>Posterior: optic tract | Anterior: lateral olfactory area<br>Middle: putamen, anterior commissure<br>Posterior: amygdalostratial transition area |

All anatomic nomenclature, abbreviations, and atlas plates are from Ding, S.L. et al. (2016). Comprehensive cellular-resolution atlas of the adult human brain. *J Comp Neurol* **524** (16): 3127-3481. Anatomic abbreviations are provided in the Supplementary Material.

**Table S7.** Interspecimen Variance in Brainstem-Thalamic Projection Connectivity Probabilities.

| Nodes | IL |  |  | PaV |  |  | Ret |  |  |
| --- | --- | --- | --- | --- | --- | --- | --- | --- | --- |
|  | S2 | S3 | Mean | S2 | S3 | Mean | S2 | S3 | Mean |
| mRt | 0.60 | 3.05 | 1.82 | 0.4 | 3.07 | 1.72 | 1.94 | 3.96 | 2.95 |
| VTA | 0.03 | 0.70 | 0.36 | 0 | 0.75 | 0.39 | 0.09 | 0.80 | 0.44 |
| PAG | 0.25 | 3.26 | 1.75 | 1.7 | 6.33 | 4.02 | 2.14 | 0.67 | 1.40 |
| PTg | 0.02 | 0.72 | 0.37 | 0 | 0.80 | 0.40 | 0.04 | 0.47 | 0.26 |
| DR | 0.001 | 0.16 | 0.08 | 0 | 0.08 | 0.04 | 0.002 | 0.20 | 0.10 |
| PnO | 0 | 0.10 | 0.05 | 0 | 0.003 | 0.001 | 0 | 0.34 | 0.17 |
| PBC | 0 | 0.25 | 0.13 | 0 | 0.01 | 0.003 | 0 | 0.49 | 0.25 |
| MnR | 0 | 0.11 | 0.05 | 0 | 0.003 | 0.001 | 0 | 0.3 | 0.15 |
| LC | 0 | 0.10 | 0.05 | 0 | 0.003 | 0.002 | 0 | 0.1 | 0.05 |
| LDTg | 0 | 0.07 | 0.04 | 0 | 0.002 | 0.001 | 0 | 0.10 | 0.05 |

All connectivity probability measurements are multiplied by 100 and reported as percentages. Anatomic abbreviations are provided in the Supplementary Material. Highlighted in RED are connections that failed that confidence interval test. S2 = Specimen 2; S3 = Specimen 3.

**Table S8.** Interspecimen Variance in Brainstem-Hypothalamic Projection Connectivity Probabilities.

| Nodes | TM |  |  | LHA |  |  | SUM |  |  |
| --- | --- | --- | --- | --- | --- | --- | --- | --- | --- |
|  | S2 | S3 | Mean | S2 | S3 | Mean | S2 | S3 | Mean |
| mRt | 1.15 | 1.47 | 1.31 | 3.79 | 3.48 | 3.64 | 0.76 | 0.93 | 0.85 |
| VTA | 10.20 | 18.71 | 14.45 | 22.57 | 24.29 | 23.43 | 7.85 | 23.48 | 15.66 |
| PAG | 0.55 | 2.82 | 1.69 | 3.02 | 2.74 | 2.88 | 0.43 | 2.39 | 1.41 |
| PTg | 9.96 | 9.57 | 9.77 | 9.98 | 13.48 | 11.73 | 5.89 | 8.88 | 7.36 |
| DR | 1.49 | 3.76 | 2.62 | 2.80 | 2.79 | 2.79 | 1.43 | 4.47 | 2.95 |
| PnO | 1.84 | 1.70 | 1.77 | 2.57 | 1.61 | 2.09 | 1.37 | 2.85 | 2.11 |
| PBC | 3.99 | 4.05 | 4.02 | 4.20 | 4.73 | 4.46 | 3.17 | 3.77 | 3.47 |
| MnR | 0.94 | 1.01 | 0.98 | 1.07 | 0.71 | 0.89 | 1.10 | 2.01 | 1.55 |
| LC | 0.88 | 1.91 | 1.39 | 1.45 | 2.81 | 2.13 | 1.05 | 2.44 | 1.74 |
| LDTg | 0.36 | 1.36 | 0.86 | 0.75 | 1.96 | 1.36 | 0.56 | 2.26 | 1.41 |

All connectivity probability measurements are multiplied by 100 and reported as percentages. Anatomic abbreviations are provided in the Supplementary Material. Highlighted in RED are connections that failed that confidence interval test. S2 = Specimen 2; S3 = Specimen 3.

**Table S9.** Interspecimen Variance in Brainstem-Basal Forebrain Projection Connectivity Probabilities.

| Nodes | DBB |  |  | BNM/SI |  |  |
| --- | --- | --- | --- | --- | --- | --- |
|  | S2 | S3 | Mean | S2 | S3 | Mean |
| **mRt** | 0.03 | 0.07 | 0.05 | 1.23 | 1.52 | 1.37 |
| **VTA** | 0.08 | 0.006 | 0.04 | 6.61 | 10.93 | 8.77 |
| **PAG** | 0.07 | 0.06 | 0.07 | 1.01 | 0.92 | 0.96 |
| **PTg** | 0.01 | 0.01 | 0.01 | 1.91 | 6.78 | 4.35 |
| **DR** | 0.04 | 0.04 | 0.04 | 0.80 | 1.12 | 0.96 |
| **PnO** | 0.003 | 0.001 | 0.002 | 0.71 | 0.72 | 0.71 |
| **PBC** | 0.009 | 0.006 | 0.007 | 0.95 | 2.22 | 1.58 |
| **MnR** | 0.002 | 0.001 | 0.001 | 0.29 | 0.35 | 0.32 |
| **LC** | 0.01 | 0.02 | 0.01 | 0.42 | 1.32 | 0.87 |
| **LDtg** | 0.005 | 0.006 | 0.006 | 0.22 | 0.97 | 0.59 |

All connectivity probability measurements are multiplied by 100 and reported as percentages. Anatomic abbreviations are provided in the Supplementary Material. Highlighted in **RED** are connections that failed that confidence interval test. S2 = Specimen 2; S3 = Specimen 3.

**Table S10.**
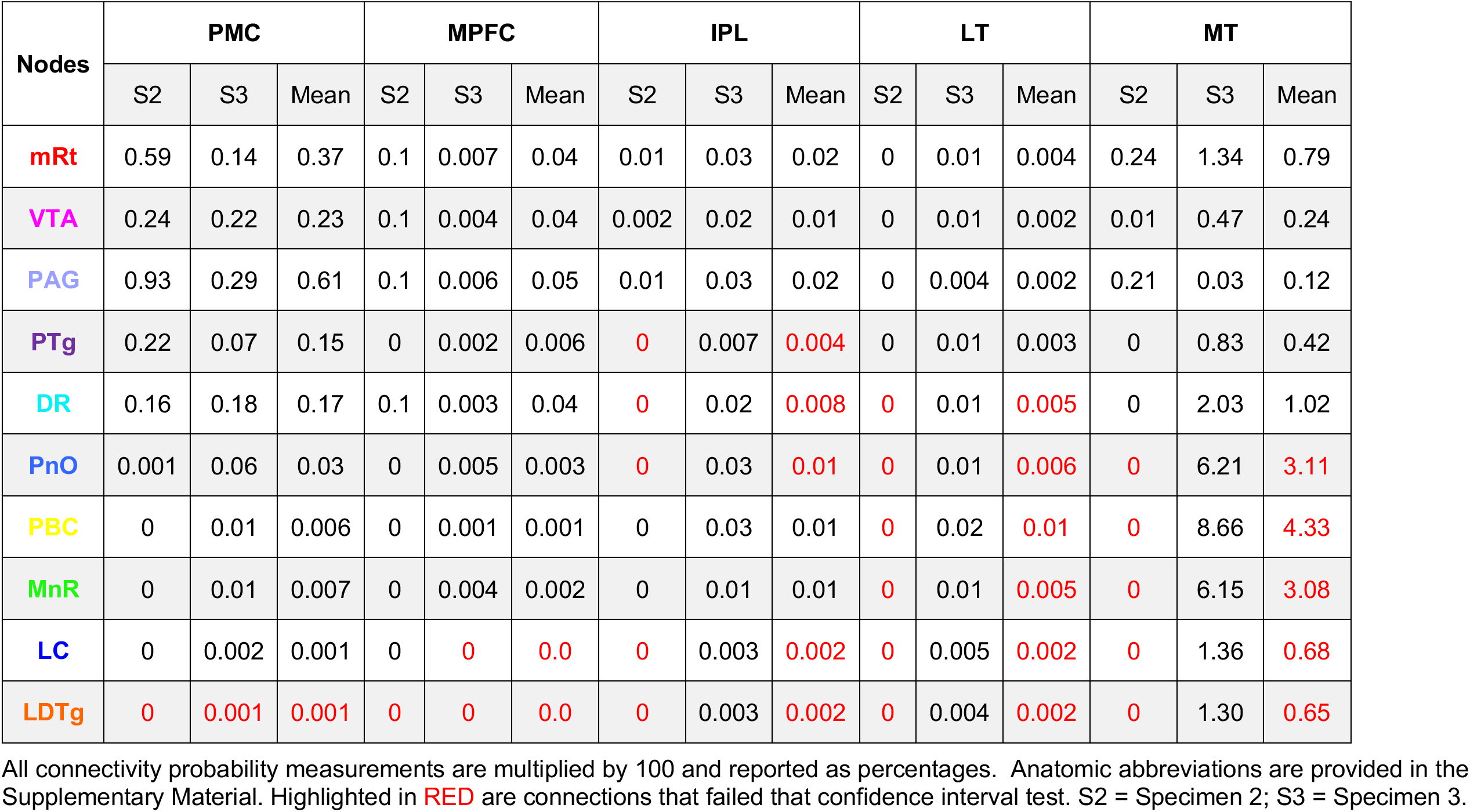
Interspecimen Variance in Brainstem-DMN Projection Connectivity Probabilities.

**Table S11.** Interspecimen Variance in Left-Sided Association Connectivity Probabilities.

| Nodes | mRt_L |  |  | PTg_L |  |  | PnO_L |  |  | PBC_L |  |  | LC_L |  |  | LDTg_L |  |  |
| --- | --- | --- | --- | --- | --- | --- | --- | --- | --- | --- | --- | --- | --- | --- | --- | --- | --- | --- |
|  | S2 | S3 | Mean | S2 | S3 | Mean | S2 | S3 | Mean | S2 | S3 | Mean | S2 | S3 | Mean | S2 | S3 | Mean |
| mRt_L |  |  |  | 23.00 | 19.36 | 21.01 | 0.42 | 0.01 | 0.21 | 0.35 | 0.03 | 0.19 | 0.10 | 0.72 | 0.41 | 0.06 | 0.11 | 0.08 |
| PTg_L | 22.67 | 19.36 | 21.01 |  |  |  | 1.57 | 0.05 | 0.81 | 1.53 | 3.47 | 2.50 | 0.93 | 0.04 | 0.49 | 0.46 | 0.02 | 0.24 |
| PnO_L | 0.42 | 0.01 | 0.21 | 1.60 | 0.05 | 0.81 |  |  |  | 27.60 | 40.93 | 34.26 | 18.47 | 3.34 | 10.90 | 14.40 | 1.93 | 8.16 |
| PBC_L | 0.35 | 0.03 | 0.19 | 1.50 | 3.47 | 2.50 | 27.60 | 40.93 | 34.26 |  |  |  | 17.57 | 30.81 | 24.19 | 6.21 | 28.30 | 17.26 |
| LC_L | 0.10 | 0.72 | 0.41 | 0.90 | 0.04 | 0.49 | 18.47 | 3.34 | 10.90 | 17.57 | 30.81 | 24.19 |  |  |  | 52.00 | 58.09 | 55.05 |
| LDTg_L | 0.06 | 0.11 | 0.08 | 0.50 | 0.02 | 0.24 | 14.40 | 1.93 | 8.16 | 6.21 | 28.30 | 17.26 | 52.00 | 58.09 | 55.05 |  |  |  |

All connectivity probability measurements are multiplied by 100 and reported as percentages. Anatomic abbreviations are provided in the Supplementary Material. Highlighted in **RED** are connections that failed that confidence interval test. S2 = Specimen 2; S3 = Specimen 3.

**Table S12.** Interspecimen Variance in Right-Sided Association Connectivity Probabilities.

| Nodes | mRt_R |  |  | PTg_R |  |  | PnO_R |  |  | PBC_R |  |  | LC_R |  |  | LDTg_R |  |  |
| --- | --- | --- | --- | --- | --- | --- | --- | --- | --- | --- | --- | --- | --- | --- | --- | --- | --- | --- |
|  | S2 | S3 | Mean | S2 | S3 | Mean | S2 | S3 | Mean | S2 | S3 | Mean | S2 | S3 | Mean | S2 | S3 | Mean |
| mRt_R |  |  |  | 7.52 | 21.34 | 14.43 | 3.35 | 0.17 | 1.76 | 24.88 | 17.93 | 21.41 | 28.29 | 27.89 | 28.09 | 11.23 | 5.28 | 8.25 |
| PTg_R | 7.52 | 21.34 | 14.43 |  |  |  | 2.81 | 0.75 | 1.78 | 45.81 | 58.66 | 52.23 | 2.12 | 3.41 | 2.76 | 0.65 | 0.17 | 0.41 |
| PnO_R | 3.35 | 0.17 | 1.76 | 2.81 | 0.75 | 1.78 |  |  |  | 2.94 | 2.44 | 2.69 | 1.43 | 3.43 | 2.43 | 2.21 | 1.65 | 1.93 |
| PBC_R | 24.88 | 17.93 | 21.41 | 45.81 | 58.66 | 52.23 | 2.94 | 2.44 | 2.69 |  |  |  | 17.86 | 6.80 | 12.33 | 0.68 | 0.26 | 0.47 |
| LC_R | 28.29 | 27.89 | 28.09 | 2.12 | 3.41 | 2.76 | 1.431 | 3.43 | 2.43 | 17.86 | 6.80 | 12.33 |  |  |  | 50.79 | 31.60 | 41.19 |
| LDTg_R | 11.23 | 5.28 | 8.25 | 0.65 | 0.17 | 0.41 | 2.21 | 1.65 | 1.93 | 0.68 | 0.26 | 0.47 | 50.79 | 31.60 | 41.19 |  |  |  |

All connectivity probability measurements are multiplied by 100 and reported as percentages. Anatomic abbreviations are provided in the Supplementary Material. Highlighted in **RED** are connections that failed that confidence interval test. S2 = Specimen 2; S3 = Specimen 3.

**Table S13.** Interspecimen Variance in Midline Association Connectivity Probabilities.

| Nodes | VTA |  |  | PAG |  |  | DR |  |  | MnR |  |  |
| --- | --- | --- | --- | --- | --- | --- | --- | --- | --- | --- | --- | --- |
|  | S2 | S3 | Mean | S2 | S3 | Mean | S2 | S3 | Mean | S2 | S3 | Mean |
| mRt_L | 16.29 | 13.77 | 15.03 | 50.00 | 9.37 | 29.83 | 10.20 | 3.99 | 7.09 | 0.66 | 0.01 | 0.33 |
| PTg_L | 3.72 | 1.95 | 2.84 | 4.60 | 0.62 | 2.63 | 2.19 | 0.07 | 1.13 | 2.55 | 0.02 | 1.29 |
| PnO_L | 4.49 | 0.02 | 2.25 | 0.10 | 0.01 | 0.07 | 8.66 | 6.51 | 7.58 | 62.61 | 68.11 | 65.36 |
| PBC_L | 0.10 | 0.03 | 0.07 | 0 | 0.01 | 0.01 | 3.38 | 20.34 | 11.86 | 22.09 | 36.38 | 29.23 |
| LC_L | 0.25 | 0.02 | 0.13 | 0 | 0.03 | 0.02 | 8.85 | 18.12 | 13.49 | 25.24 | 5.98 | 15.61 |
| LDTg_L | 0.26 | 0.01 | 0.13 | 0 | 0.01 | 0.01 | 6.56 | 12.25 | 9.41 | 21.65 | 4.60 | 13.13 |
| mRt_R | 9.42 | 13.86 | 11.64 | 50.00 | 20.35 | 35.29 | 10.00 | 5.79 | 7.90 | 0.14 | 0.01 | 0.07 |
| PTg_R | 7.89 | 8.06 | 7.98 | 1.10 | 8.75 | 4.91 | 1.25 | 2.80 | 2.02 | 0.25 | 0.02 | 0.14 |
| PnO_R | 6.06 | 0.01 | 3.04 | 0.50 | 0.01 | 0.27 | 19.05 | 7.48 | 13.26 | 64.63 | 64.02 | 64.32 |
| PBC_R | 2.58 | 0.21 | 1.40 | 0.50 | 0.45 | 0.47 | 8.42 | 15.38 | 11.90 | 7.33 | 49.39 | 28.36 |
| LC_R | 1.30 | 0.12 | 0.71 | 0.30 | 0.02 | 0.14 | 14.03 | 18.80 | 16.42 | 8.81 | 0.91 | 4.86 |
| LDTg_R | 0.58 | 0.01 | 0.30 | 0.10 | 0.01 | 0.04 | 8.42 | 15.82 | 12.12 | 4.64 | 1.31 | 2.98 |
| VTA |  |  |  | 14.06 | 34.95 | 24.51 | 27.28 | 17.34 | 22.31 | 6.69 | 0.01 | 3.35 |
| PAG | 14.06 | 34.95 | 24.51 |  |  |  | 26.83 | 26.44 | 26.63 | 0.27 | 0.01 | 0.14 |
| DR | 27.28 | 17.34 | 22.31 | 26.83 | 26.44 | 26.63 |  |  |  | 19.63 | 13.34 | 16.49 |
| MnR | 6.69 | 0.01 | 3.35 | 0.27 | 0.01 | 0.14 | 19.63 | 13.34 | 16.49 |  |  |  |

All connectivity probability measurements are multiplied by 100 and reported as percentages. Anatomic abbreviations are provided in the Supplementary Material. Highlighted in **RED** are connections that failed that confidence interval test. S2 = Specimen 2; S3 = Specimen 3.

**Table S14.** Interspecimen Variance in Commissural Connectivity Probabilities.

| Nodes | mRt_R |  |  | PTg_R |  |  | PnO_R |  |  | PBC_R |  |  | LC_R |  |  | LDTg_R |  |  |
| --- | --- | --- | --- | --- | --- | --- | --- | --- | --- | --- | --- | --- | --- | --- | --- | --- | --- | --- |
|  | S2 | S3 | Mean | S2 | S3 | Mean | S2 | S3 | Mean | S2 | S3 | Mean | S2 | S3 | Mean | S2 | S3 | Mean |
| mRt_L | 35.25 | 0.21 | 17.73 | 0.70 | 0.11 | 0.41 | 0.78 | 0.01 | 0.40 | 0.17 | 0.01 | 0.09 | 0.32 | 0.004 | 0.16 | 0.05 | 0.003 | 0.02 |
| PTg_L | 4.28 | 0.13 | 2.20 | 3.90 | 0.14 | 2.03 | 2.29 | 0.02 | 1.16 | 0.62 | 0.03 | 0.33 | 1.46 | 0.01 | 0.74 | 0.20 | 0.02 | 0.11 |
| PnO_L | 0.04 | 0.02 | 0.03 | 0.10 | 0.04 | 0.06 | 33.46 | 59.52 | 46.49 | 3.23 | 49.19 | 26.21 | 4.45 | 1.02 | 2.73 | 2.33 | 1.25 | 1.79 |
| PBC_L | 0.01 | 0.01 | 0.01 | 0 | 0.06 | 0.03 | 9.45 | 31.87 | 20.66 | 0.10 | 48.70 | 24.40 | 0.55 | 25.48 | 13.01 | 0.13 | 27.30 | 13.72 |
| LC_L | 0.01 | 0.01 | 0.01 | 0 | 0.03 | 0.02 | 20.60 | 2.56 | 11.58 | 0.38 | 23.72 | 12.05 | 3.35 | 30.35 | 16.85 | 1.02 | 38.11 | 19.56 |
| LDTg_L | 0.01 | 0.002 | 0.004 | 0 | 0.01 | 0.01 | 18.68 | 2.03 | 10.35 | 0.49 | 17.32 | 8.91 | 3.47 | 25.04 | 14.26 | 2.25 | 43.88 | 23.07 |

All connectivity probability measurements are multiplied by 100 and reported as percentages. Anatomic abbreviations are provided in the Supplementary Material. Highlighted in RED are connections that failed that confidence interval test. S2 = Specimen 2; S3 = Specimen 3.

## Acknowledgements

We dedicate this work in memoriam to Marty Samuels, MD, the Miriam Sydney Joseph Distinguished Professor of Neurology at Harvard Medical School and Founding Chair of the Department of Neurology at Brigham and Women’s Hospital. Dr. Samuels mentored generations of neurologists, neuroimagers, and neuropathologists, of which we were fortunate to be a part. Dr. Samuels was an avid and insightful student and teacher of neuroanatomy and neuropathology of human consciousness, from whom we learned much. He was an early supporter of the histologic-radiology correlation studies performed here, for which we are grateful.

We are grateful to the families for the brain donation of their loved ones, and to the individuals who donated their own brains for this study to advance medical science. We also thank Bruce Rosen, Merit Cudkowicz, and Jonathan Rosand for their support of this work. We thank Michelle Siciliano for assistance with processing of human brain specimens and Kathryn Regan at Horus Scientific for assistance with immunostaining. We thank Kimberly Main Knoper for artwork used in the connectograms.

## Author contributions

B.L.E., R.D.F., B.F., and H.C.K. conceived, designed, and coordinated the study and acquired funding. A.v.d.K. and B.F. developed the ex vivo diffusion MRI protocol for data acquisition. M.O., H.J.F., S.B.S., J.E.I., L.Z., Y.G.B., J.A., D.N.G., B.R.D., C.M., A.S., developed the ex vivo diffusion MRI processing and analysis methods. J.A., R.L.H., R.D.F., and H.C.K. developed the histology and immunostaining protocols. B.L.E., M.O., J.A., R.L.H., R.D.F., and H.C.K., interpreting the histology and immunostaining data. B.L.E. and M.O. manually traced the nodal regions of interest on the ex vivo diffusion MRI datasets, with anatomic confirmation performed by H.C.K. The statistical plan was developed by E.N.B. J.L. developed the methods for the in vivo functional MRI (fMRI) connectivity analysis and processed and analyzed the fMRI data. Connectivity data were interpreted by B.L.E., M.O., H.J.F., J.L., C.M., S.B.S., J.E.I., L.Z., J.A., Y.G.B., R.L.H., D.N.G., J.T.G., C.D., E.N.B., R.D.F., B.F., H.C.K. B.L.E. and H.C.K. drafted the manuscript. All authors provided critical commentary and revisions and reviewed the final manuscript.

## Data and materials availability

All ex vivo diffusion MRI data, along with the node regions of interest used for the connectivity analyses, are available at (https://openneuro.org/datasets/ds004640/versions/1.0.1). All histology and immunostain data are available at (https://histopath.nmr.mgh.harvard.edu). Version 2.0 of the Harvard Ascending Arousal Network Atlas is available at Dryad Data Repository (doi:10.5061/dryad.zw3r228d2; download link: https://datadryad.org/stash/share/aJ713eXY12ND56bzOBejVG2jmOFCD2CKxdSJsYWEHkw). The minimally preprocessed in vivo fMRI data are available athttps://db.humanconnectome.org, and our processed fMRI data are available at https://openneuro.org/datasets/ds003716/versions/1.0.0. All code used to process the ex vivo MRI data are available at https://github.com/ComaRecoveryLab/Network-based_Autopsy, and all code used to process the in vivo fMRI data are available at https://github.com/ComaRecoveryLab/Subcortical_DMN_Functional_Connectivity.

